# Explainable Deep Learning Framework: Decoding Brain Task and Prediction of Individual Performance in False-Belief Task at Early Childhood Stage

**DOI:** 10.1101/2024.02.29.582682

**Authors:** Km Bhavna, Azman Akhter, Romi Banerjee, Dipanjan Roy

## Abstract

Decoding of brain tasks aims to identify individuals’ brain states and brain fingerprints to predict behavior. Deep learning provides an important platform for analyzing brain signals at different developmental stages to understand brain dynamics. Due to their internal architecture and feature extraction techniques, existing machine learning and deep-learning approaches for fMRI-based brain decoding must improve classification performance and explainability. The existing approaches also focus on something other than the behavioral traits that can tell about individuals’ variability in behavioral traits. In the current study, we hypothesized that even at the early childhood stage (as early as 3 years), connectivity between brain regions could decode brain tasks and predict behavioural performance in false-belief tasks. To this end, we proposed an explainable deep learning framework to decode brain states (Theory of Mind and Pain states) and predict individual performance on ToM-related false-belief tasks in a developmental dataset. We proposed an explainable spatiotemporal connectivity-based Graph Convolutional Neural Network (Ex-stGCNN) model for decoding brain tasks. Here, we consider a dataset (age range: 3-12 yrs and adults, samples: 155) in which participants were watching a short, soundless animated movie, ”Partly Cloudy,” that activated Theory-of-Mind (ToM) and pain networks. After scanning, the participants underwent a ToM-related false-belief task, leading to categorization into the pass, fail, and inconsistent groups based on performance. We trained our proposed model using Static Functional Connectivity (SFC) and Inter-Subject Functional Correlations (ISFC) matrices separately. We observed that the stimulus-driven feature set (ISFC) could capture ToM and Pain brain states more accurately with an average accuracy of 94%, whereas it achieved 85% accuracy using SFC matrices. We also validated our results using five-fold cross-validation and achieved an average accuracy of 92%. Besides this study, we applied the SHAP approach to identify neurobiological brain fingerprints that contributed the most to predictions. We hypothesized that ToM network brain connectivity could predict individual performance on false-belief tasks. We proposed an Explainable Convolutional Variational Auto-Encoder model using functional connectivity (FC) to predict individual performance on false-belief tasks and achieved 90% accuracy.

## 1 Introduction

Decoding of behavioral tasks and cognitive states from the brain activity, or simply the ’brain decoding’ has emerged as one of the most active research areas because of its potentially wide-ranging implications in medical and therapeutic engineering fields [Hou et al., 2022, Santhanam et al., 2006]. However, successfully building models that can generalize over different cognitive states and tasks, including higher order and complex cognitive functions like social cognition, from early childhood and adolescence (developmental phase) to the point of maturation of behavioral functions, using only short-duration brain recordings that too from a small set of subjects, is still a huge challenge. Functional magnetic resonance imaging (fMRI) has been widely used in such brain decoding applications due to its spatial and temporal resolution and noninvasive nature. Traditional fMRI techniques, employing a general linear model, infer that if a subject performs some task or is in some cognitive state, then particular brain regions will be activated. This direct inference is sometimes misinterpreted and stated in the form of its reverse inference that if some particular pattern of brain region(s) are activated, then the participants would be experiencing some specific cognitive state [Poldrack, 2011,Zhang et al., 2021]. However, this presumption is also not always accurate because certain brain areas activated by various tasks might correspond with one another. Another notable limitation is that not all behavioral tasks can engage certain parts of the brain to a considerable activation range. The activated regions, detected, rely heavily on the nature of the behavior and also on the sample size. In the literature [Zhang et al., 2021, Poldrack, 2006], authors argue that reverse inference can be accurately implemented using brain decoding techniques. Using the brain decoding technique, a certain spatio-temporal pattern of brain activity is predictive of a particular cognitive state. Brain decoding could be performed with a large number of experimental conditions. As [Poldrack, 2011] mentions ”decoding should generalize over a large and diverse class of experimental conditions, to ensure that the association between a pattern of brain activity and a cognitive state reflects a key property of the corresponding regions and not a mere reflection of the context where this association was tested.

In the field of brain (or neural) decoding, brain activities are recorded during various cognitive states, evoked using a variety of stimuli (or experimental conditions) to interpret the associated information [Zhang et al., 2021, Haxby et al., 2001, Haxby et al., 2014, Huth et al., 2012, Wang et al., 2020]. BOLD signals from fMRI data have been used to decode cognitive states associated with not just visual stimuli, but audio stimuli [Song et al., 2013], and beyond perception-level processes and tasks as well, such as language [Correia et al., 2014], and working memory [Spellman et al., 2015]. Some studies [Pilgramm et al., 2016] have also shown that motor imagery can be decoded from spatial patterns of fMRI signals. There are well-known ML based methods like MultiVoxels Pattern Analysis (MVPA) as well, used to identify affective states, valence, and arousal [Baucom et al., 2012]. Machine learning and deep learning models have been successful in brain decoding tasks [Zhang et al., 2021, Zhang and Bellec, 2019, Hou et al., 2022, Ye et al., 2022]. Although previous studies [Haxby et al., 2001, Li and Fan, 2019, Wang et al., 2020] have demonstrated significant improvements in neural decoding about specific cognitive or behavioral tasks, few have attempted to decode various functions across a wide range of behavioral domains.

According to Zang et al. [Zhang et al., 2021], Meta-analytic methodologies have been utilized for multi-domain brain decoding [Bartley et al., 2018]. However, meta-analyses have several additional limitations, including inconsistent samples across various cognitive domains, publication bias towards positive effects, and overestimated effect sizes derived from small-sampled studies [Alamolhoda et al., 2017, Zhang et al., 2021, Dubben and Beck-Bornholdt, 2005, Lin, 2018]. Given the few available research, these measures may introduce bias into the decoding analysis by leading to incorrect inferences about the cognitive states [Lieberman et al., 2016, Lieberman and Eisenberger, 2015, Wager et al., 2016]. An alternate method is to overcome these biases by training linear classifiers on activation maps acquired from a group of individuals who have been scanned. At the same time, they performed several different cognitive activities [Zhang et al., 2021, Bzdok et al., 2016, Poldrack et al., 2009, Varoquaux et al., 2018]. Deep learning models such as Convolutional Neural Networks (CNN) are efficient and scalable, that can differentiate patterns in various tasks without requiring manual features. 3d CNN-based models were able to perform decoding of brain tasks based on the Human Connectome Project (HCP) dataset, a multidomain fMRI dataset. [Wang et al., 2020]. Firstly, it was not investigated whether those CNNs would generalize to the data from the developmental period (early childhood and pre-adolescence). Secondly, though CNN performs admirably with grid-like inputs in Euclidean space, such as natural images, there is a possibility that it is not suited for use with data residing in non-Euclidean space, nonetheless, it is known that the neuronal activity between many different brain areas may be strongly synchronized, giving rise to certain active brain networks like Default-Mode Networks (DMN) during the resting state, and during other cognitive paradigms [Poldrack, 2006]. As a result, the geometric distance in Euclidean space may not adequately represent the functional distance between different parts of the brain (see similar with LFP recordings from Monkey’s Visual cortex [Rosenbaum et al., 2017]). Instead, some geometric deep learning methods, such as graph convolutional networks (GCNs), would better suit non-Euclidean data types, such as graphs [Zhang et al., 2021, Zhang and Bellec, 2019].

Most studies above utilize controlled task paradigms or resting-state data that fail to capture how the brain typically functions and responds in ecologically valid contexts. That raises concerns about the replicability and usefulness of the previous evidence for real-life scenarios. Naturalistic stimuli provide a promising pathway to examine brain dynamics across a rich and diverse rational spectrum of realistic human experience(s) [Simony and Chang, 2020]. Moreover, naturalistic paradigms provide a rich context-dependent array of cognitive states and sub-states to investigate with the help of ML tools. However, what specific measures to be used that could capture a robust correlation between the recorded fMRI signals and cognitive states that emerge in response to realistic stimuli for developmental datasets or otherwise still needs to be properly established. Previous studies [Ye et al., 2023, Zhang et al., 2021] have shown decoding of brain tasks in adolescence and adult data, whereas decoding brain tasks in early childhood (as early as 3 years) and pre-adolescence is challenging, we believe, primarily due to limitations of deep neural networks (DNN) models, in the inclusion of established brain-inspired architecture, such as lateral connections within a neural-network layer and extensive connectivity between layers. Last but not least of the issues in our understanding, inter-subject variability is also challenging when dealing with developmental data.

Given the above account, In this study, we decided to tackle mainly the following issues: 1) Brain connectivity-based decoding of stimulus-driven states, 2) Decoding brain activity across cognitive states specifically for high-order cognition such as mentalization and pain processing tasks, 3) Decoding and scene predictions using short time-course data from single fMRI session and small sample size, 4) Integrating brain-like non-Euclidian, i.e., graph-based modeling approach, and 5) decoding brain activity evoked by naturalistic stimuli.

To systematically define the framework, first, we hypothesized that a spatiotemporal connectivity-based graph convolutional neural network could better capture brain states using naturalistic visual stimuli in the early childhood developmental dataset using graph laplacian approach, which models the brain as a graph in which treating region-of-interest (ROI) as nodes and their connectivity as edges. Moreover, as naturalistic stimuli are time-locked, we hypothesized that stimulus-driven measures could be a better feature-set to train the proposed model. Second, we hypothesized that FC between specified ROIs could predict the individual-level performance of the false-belief task. To test our hypotheses, we performed brain decoding on a naturalistic movie-watching dataset, in which participants (ages 3-12 yrs and adults (no. of samples =155)) were watching a short soundless animated movie ”Partly Cloudy” of 5.6 minutes duration that was validated to activate ToM and pain (seeing others in pain) networks. There was a single fMRI session for each subject in which some scenes evoked maximum activation in ToM and some in the pain network at some particular time points. After the scanning session, a ToM-related false-belief task was performed, and based on performance, the participants were categorized into the pass, inconsistent, and fail groups. We selected short-duration time-courses from the events of maximum activation of each of the networks of interests (total no. of time-points = 168, for details, see Results section), similar to the main study from which the data is taken [Richardson et al., 2018].

We proposed a novel Explainable spatiotemporal connectivity-based Graph Convolutional Neural Network (Ex-stGCNN) model to decode time courses where a participant was experiencing a particularly cognitive state (Refer to Figure 1). The proposed model could extract features from non-Euclidean data and process graph-structured signals. Firstly, the graph Laplacian is constructed to describe the topological connection of brain regions. This graph is constructed based on Static Functional Connectivity (SFC), which reflects inter-regional correlations arising from a mixture of stimulus-induced neural processes, intrinsic neural processes, and non-neuronal noise, and Inter-Subject Functional Connectivity (ISFC), which isolates stimulus-dependent inter-regional correlations by modeling BOLD signal of one brain on the other brain’s exposed to the same stimulus [Simony et al., 2016]. In addition, the generalised features were learned by the Ex-stGCNN when it is constructed on graph convolutional layers. The pooling layer was used for dimensionality reduction. In addition, the fully connected softmax layer was responsible for deriving the final prediction (Refer to Figure 1). Finally, the Chebyshev polynomial was used to approximate the graph convolutional filters, substantially contributing to the advancement of computing efficacy. As a result, we achieved an average of 94 % accuracy with an F1-Score of 0.95. We applied the SHAP (SHapley Additive exPlanations) method for explainability and finally identified neurobiological brain features responsible for prediction.

**Figure 1:**
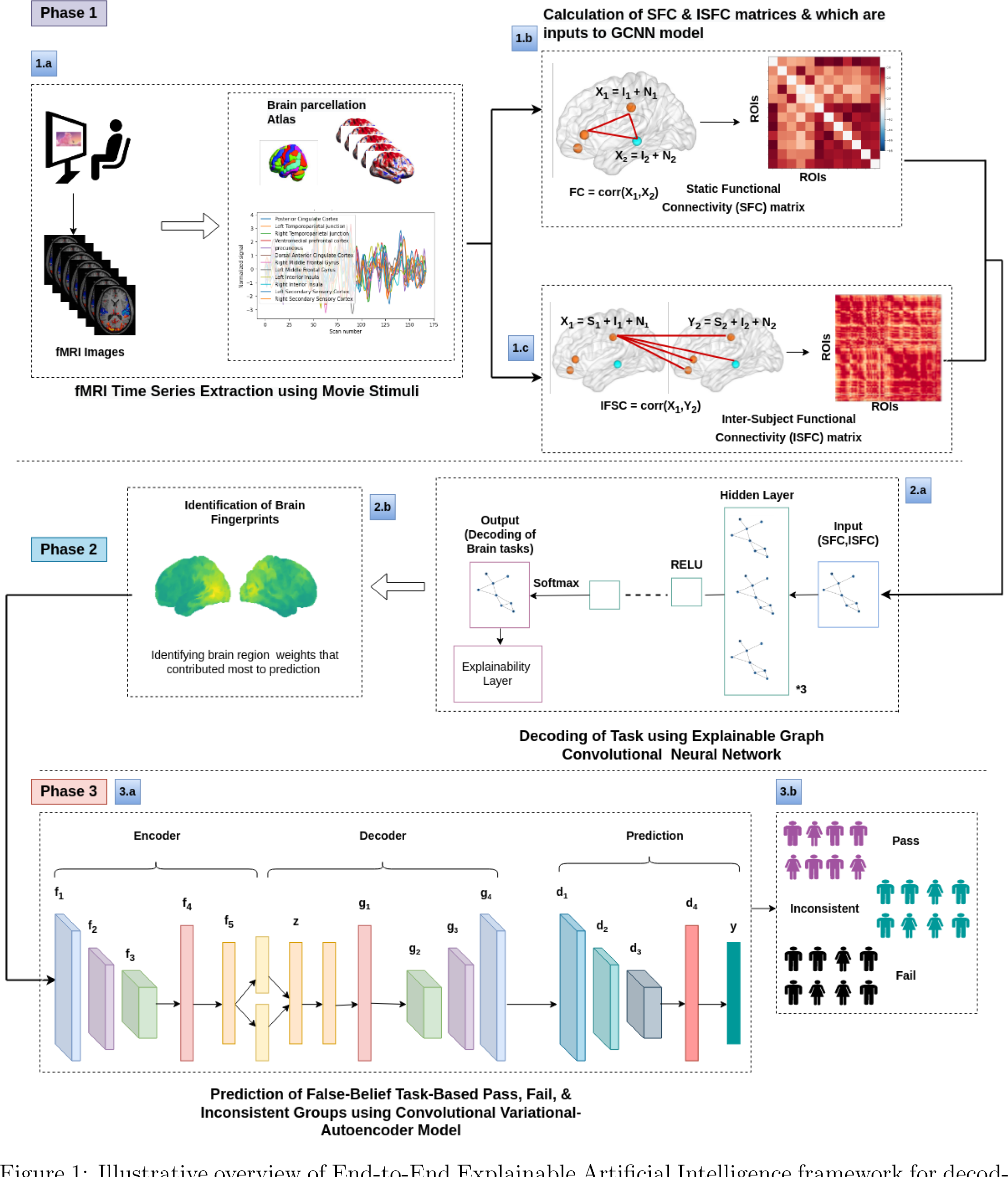
Illustrative overview of End-to-End Explainable Artificial Intelligence framework for decoding brain tasks and false-belief task-based pass, fail, and inconsistent groups. The contribution of the proposed work is as follows: **Phase 1:** 1.a) data collection during naturalistic movie watching and extraction of time-series from ToM and pain networks; 1.b) and 1.c) Calculation of SFC and ISFC matrices, **Phase 2:** 2.a) SFC and ISFC matrices are separately provided as input to Ex-stGCNN model to decode brain tasks; 2.b) Identifying brain fingerprints that contributed most to the prediction, **Phase 3:**, 3.a) Architecture of Ex-Convolutional VAE model to predict false-belief task-based pass, fail, and inconsistent groups using SFC matrices of ToM and pain networks; 3.b) Final groups were categorized into three classes: Pass (5-6 correct answers), inconsistent (3-4 correct answers), and fail (0-2 correct answers). [Jacoby et al., 2016, Reher and Sohn, 2009, Astington and Edward, 2010, Richardson et al., 2018]. A 3-Tesla Siemens Tim Trio scanner at the Athinoula A. Martinos Imaging Center at MIT collected whole-brain structural and functional MRI data [Keil et al., 2011]. Younger participants (n = 3, M(s.d.) = 3.91(.42) years) and older participants (n = 28, M(s.d.) = 4.07(.42) years) of the 32-channel phased-array head coils were customized for usage by children under the age of five years old [Richardson et al., 2018]. All other participants used the Siemens 32-channel head coil. With a factor of three for GRAPPA parallel imaging, 176 interleaved sagittal slices of 1 mm isotropic voxels were used to get T1-weighted structural images (FOV: 192 mm for child coils, 256 mm for adult coils). The whole brain was covered by 32 interleaved near-axial slices that were aligned with the anterior/posterior commissure and used a gradient-echo EPI sequence sensitive to BOLD contrast to capture functional data (EPI factor: 64; TR: 2 s, TE: 30 ms, flip angle: 90°) [Richardson et al., 2018]. All functional data were upsampled in normalized space to 2 mm isotropic voxels. Based on the participant’s head motion, one TR back, prospective acquisition correction was used to modify the gradient locations. The dataset was preprocessed using SPM 8 and other toolboxes available for Matlab [Penny et al., 2011], which registered all functional images to the first run image and then registered that image to each participant’s structural images [Astington and Edward, 2010]. All structural images were normalized to Montreal Neurological Institute (MNI) template [Cantlon et al., 2006, Burgund et al., 2002]. The smoothening for all images was performed using a Gaussian filter and identified Artifactual timepoints using ART toolbox [Whitfield-Gabrieli et al., 2011, Astington and Edward, 2010].

After identifying brain features (or regions) involved in Pain and ToM (events), which helped in highly accurate ToM and Pain event decoding, we expected that the connectivity between these brain regions could be a fingerprint to predict whether the participant would pass, fail, or be inconsistent on ToM-based false-belief tasks. We implemented the unsupervised Explainable Convolutional Variational Autoencoder model (Ex-Convolutional VAE) to predict individual performance in false-belief tasks. The Ex-Convolutional VAE model included two components: (1) an encoder, which transforms the original data space (X) into a compressed low-dimensional latent space (Z), and a decoder, which reconstructs the original data by sampling from the low-dimensional latent space. (2) Use of latent space for prediction using ADAM optimizer. We obtained 90 % accuracy using non-stimulus specific SFC matrices as a feature set with an F1-Score of 0.92. To validate the results, we implemented Five-fold cross-validation. This framework not only decoded brain states from developmental age groups and highly imbalanced datasets with high accuracy from short-time course data but also predicted individual performance in false-belief tasks to categorize participants into pass, fail, and inconsistent groups successfully.

## 2 Material and methods

### 2.1 Participants and fMRI Preprocessing

To investigate the development of Theory-of-Mind and Pain networks, we analyzed an early childhood dataset (122 childhood samples (varying in age from 3 to 12 years, M(s.d)=6.7(2.3), No. of females = 64)) that also included samples from adulthood age range (33 adult samples (A total of 155 samples)) and available on OpenfMRI database [Astington and Edward, 2010, Richardson et al., 2018, Bhavna et al., 2023b]. Participants who participated in the study were from the surrounding neighborhood and brought in a signed permission form from a parent or guardian. The approval for data collection was given by the Committee on the Use of Humans as Experimental Subjects (COUHES) at the Massachusetts Institute of Technology. In this experiment, participants watched a soundless short animated movie of 5.6 minutes named ”Partly Cloudy” (Refer to Figure 2). The stimulus showed emotional and painful stories and was validated to activate ToM and Pain networks. After scanning, six explicit ToM-related questions were administered for the false-belief task to identify the correlation between brain development and behavioural scores in ToM reasoning across a wide age range of children. Each child’s performance on the ToM-related false-belief task was assessed based on the proportion of questions answered correctly out of 24 matched items (14 prediction items and 10 explanation items). Based on the outcome of these explicit false-belief task scores, the participants

**Figure 2:**
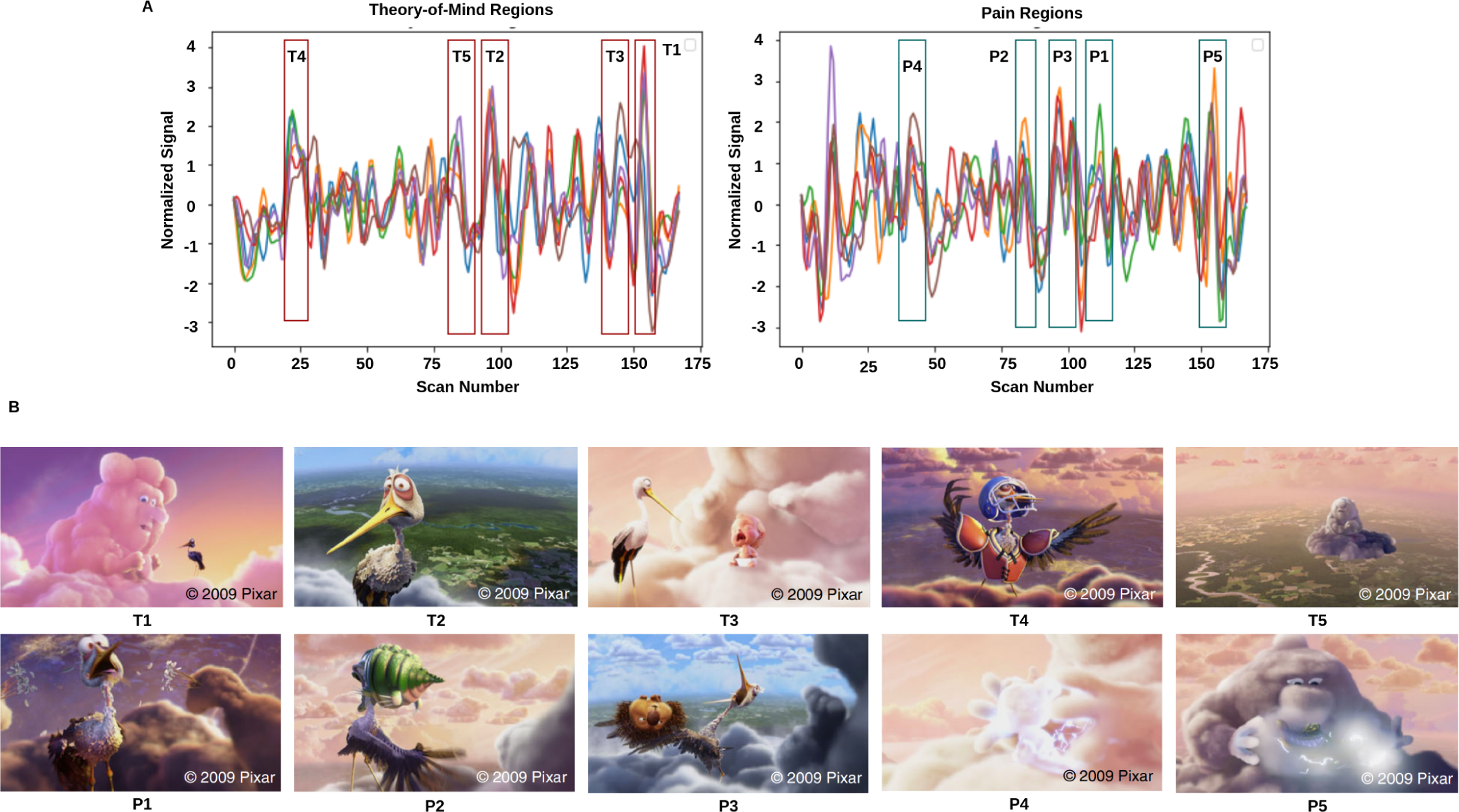
Movie demonstration: **A)** response magnitude that evoked maximum activation in ToM and pain networks. **B)** depicts the movie scenes with higher activation. *T_i_ ∈* [*T*_1_*, T*_2_*, T*_3_*, T*_4_*, T*_5_] is representing ToM scenes, and *P_i_∈* [*P*_1_*, P*_2_*, P*_3_*, P*_4_*, P*_5_] is representing pain scenes with higher activation.

### 2.2 fMRI Data Analysis and Extraction of Feature Sets

Based on the previous study, we selected six regions of interest (ROIs) from ToM network (which included bilateral Temporoparietal Junction (LTPJ and RTPJ), Posterior Cingulate Cortex (PCC), Ventro and Dorso-medial Prefrontal Cortex (vmPFC and dmPFC), and Precuneus) and six ROIs from pain network (which included bilateral Middle Frontal Gyrus (LMFG and RMFG), bilateral Interior Insula, and bilateral Secondary Sensory Cortex (LSSC and RSSC)) (Total of 12 ROIs) [Baetens et al., 2014, Mazziotta et al., 1995, Mazziotta et al., 2001]. By using MNI coordinates, we created a spherical binary mask with a 10 mm radius for all selected ROIs and extracted time-series signals (Refer to Table 1) [Richardson et al., 2018, Bhavna et al., 2023b].

**Table 1:** ToM and pain brain regions and corresponding MNI-coordinated for extracting time-series signal.

| ToM Regions |  |  | Pain Regions |  |  |
| --- | --- | --- | --- | --- | --- |
| Sr. No. | ROIs | MNI-Coordinates(X,Y,Z) | Sr. No. | ROIs | MNI-Coordinates(X,Y,Z) |
| 1 | Posterior Cingulate Cortex (PCC) | 0, -52, 18 | 1 | Right Middle Frontal Gyrus (RMFC) | 36, 38, 40 |
| 2 | Left Temporoparietal Junction (LTPJ) | -46, -68, 32 | 2 | Left Middle Frontal Gyrus (LMFC) | -36, 38, 40 |
| 3 | Right Temporoparietal Junction (RTPJ) | 46, -68, 32 | 3 | Left Interior Insula (LII) | -40, 22, 0 |
| 4 | Ventromedial Prefrontal Cortex (vmPFC) | 4, 48, -4 | 4 | Right Interior Insula (RII) | 39, 23, -4 |
| 5 | Precuneus | 0, -49, 40 | 5 | Left Secondary Sensory Cortex (LSSC) | -39, -15, 18 |
| 6 | Dorsomedial Prefrontal Cortex (dmPFC) | -10, 58, 24 | 6 | Right Secondary Sensory Cortex (RSSC) | 39, -15, 18 |

#### 2.2.1 Resting State-Functional Connectivity

To calculate functional connectivity matrices for each participant for different time courses, we calculated Pearson’s correlation using equation (1) between the average time series BOLD signals that were extracted from each of the selected brain regions [Li et al., 2017, Bhavna et al., 2023a]:

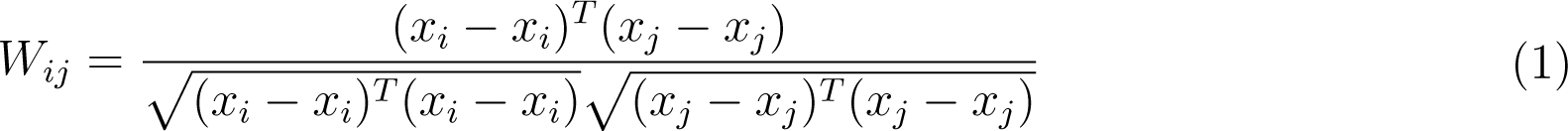

Where *x_i_∈ R^t^* is the time series related to the brain’s various areas. T is the time node, i = 1,2,…….,n, range from 1 to n, and n is the number of ROIs. Then, Fisher’s Z transformation was applied to the correlation coefficients using an inverse hyperbolic inverse function [Bhavna et al., 2023a]. In addition, we corrected each connection analysis using the false discovery rate (FDR) method.

#### 2.2.2 Computation of Inter-Subject Functional Correlations

ISFC has been used to characterize brain responses related to dynamic naturalistic cognition in a model-free way [Kim et al., 2018,Simony et al., 2016,Lynch et al., 2018, Demirtaş et al., 2019]. ISFC assesses the region-to-region neuronal coupling between subjects instead of intra-subject functional connectivity (FC), which measures the coupling inside a single sample [Hasson et al., 2004, Nastase et al., 2019]. ISFC delineates functional connectivity patterns driven by extrinsic time-locked dynamic stimuli [Hasson et al., 2004, Simony et al., 2016, Xie and Redcay, 2022]. We calculated ISFC to check the coupling between ROIs across all the subjects using equation (2) [Xie and Redcay, 2022, Kumar et al., 2020]:

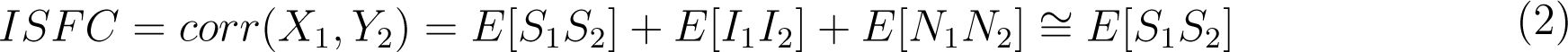

Where S was indicating task-evoked brain activity, N represented noise, and I represented intrinsic brain activity [Xie and Redcay, 2022].

### 2.3 Decoding of Task using Explainable Spatiotemporal Connectivity based Graph Convolutional Neural Network

We hypothesized that stimulus-driven brain features, ISFC, could decode brain states (ToM and Pain) more accurately than SFC features. To check our hypothesis, we implemented the Explainable Spatiotemporal connectivity-based Graph Convolutional Neural Network (Ex-stGCNN) approach to classify tasks evoked during watching stimuli. In previous work [Richardson et al., 2018], the author applied reverse correlation analysis to average response time series to determine points of maximum activation in ToM and pain networks. We accordingly selected five-time courses (*>*8 sec), from each ROI, of maximum activation in ToM and Pain networks (total of ten-time courses) (Refer to Figure 2 and Table 2). Then, we extracted time-series and converted it into a 2D matrix *T ∗ N* format for each individual where T = no. of time steps, and N = no. of regions. We calculated static functional connectivity matrices of size 12 *∗* 12 for each time course (10 matrices for each individual) [Richardson et al., 2018]. For each time course mentioned in Table 2, we also calculated ISFC matrices separately. Finally, we trained our Ex-stGCNN model in two different ways: a) using static-functional connectivity matrices and b) using ISFC matrices.

**Table 2:** Description of movie-clip events with higher activation for ToM and pain networks.

| ToM Clips Description |  | Pain Clips Description |  |
| --- | --- | --- | --- |
| Sr. No. | Description | Sr. No. | Description |
| T1 | Peck flies away to happy cloud | P1 | Gus pulls porcupine spines from Peck's head |
| T2 | Peck caught gazing at happy clouds | P2 | Alligator biting Peck |
| T3 | Baby crying, then happy | P3 | Peck tossing porcupine |
| T4 | Peck dons gear to show why he left | P4 | Cloud makes animals (lightning) |
| T5 | Pan from happy clouds to lonely cloud (Gus) | P5 | Gus makes alligator (lightning) |

#### 2.3.1 Proposed Architecture

Using PyTorch and PyTorch Geometric, the proposed model was developed in which, for every node, the Scalable Hypothesis tests (tsfresh) algorithm was used for statistical feature extraction [Kipf and Welling, 2016, Saeidi et al., 2022, Paszke et al., 2019, Fey and Lenssen, 2019]. Using the FRESH algorithm concept [Christ et al., 2016], the tsfresh algorithm combined the elements from the hypothesis tests with the feature statistical significance testing. By quantifying p-values, each created feature vector was separately analyzed to determine its relevance for the specified goal. Finally, the Benjamini-Yekutieli process determined which characteristics to preserve [Benjamini and Yekutieli,

2001]. Using node embedding methods, the high-level features associated with node were extracted. We implemented Walklets and Node2Vec node embedding algorithms to observe node attributes from graph [Perozzi et al., 2017, Grover and Leskovec, 2016]. Three convolutional layers were used in the proposed Ex-stGCNN model, where every layer had 300 neurons. The Rectified Linear Unit (ReLU) and batch normalization layers were implemented between each CNN layer to speed convergence and boost stability (Refer to Figure 3). After each CNN layer, dropout layers were applied to decrease the inherent unneeded complexity and redundant computation of the proposed multilayer Ex-stGCNN model. The final graph representation vector was calculated after applying a global mean pooling layer (Refer to Figure 3 and Table 3).

**Figure 3:**
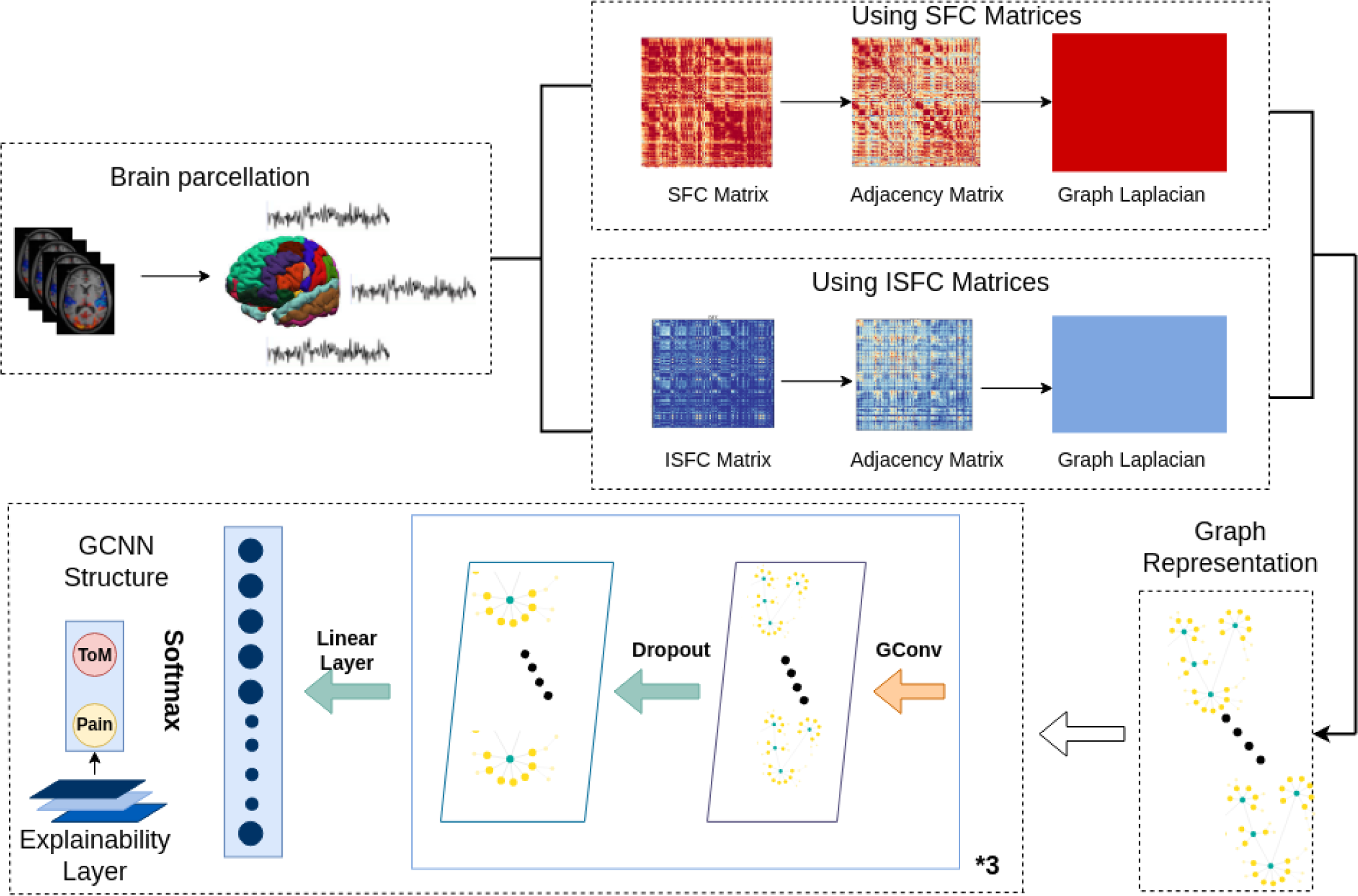
The architecture of proposed Ex-stGCNN model. First, SFC and ISFC matrices are converted into adjacency matrices and then graph Laplacian. Finally, the graph representation of these matrices is provided as input to the model for training. We trained this proposed model using SFC and ISFC matrices separately. The SHAP approach is used to extract neurological brain fingerprints to check the contribution of each brain region in the prediction.

**Table 3:**
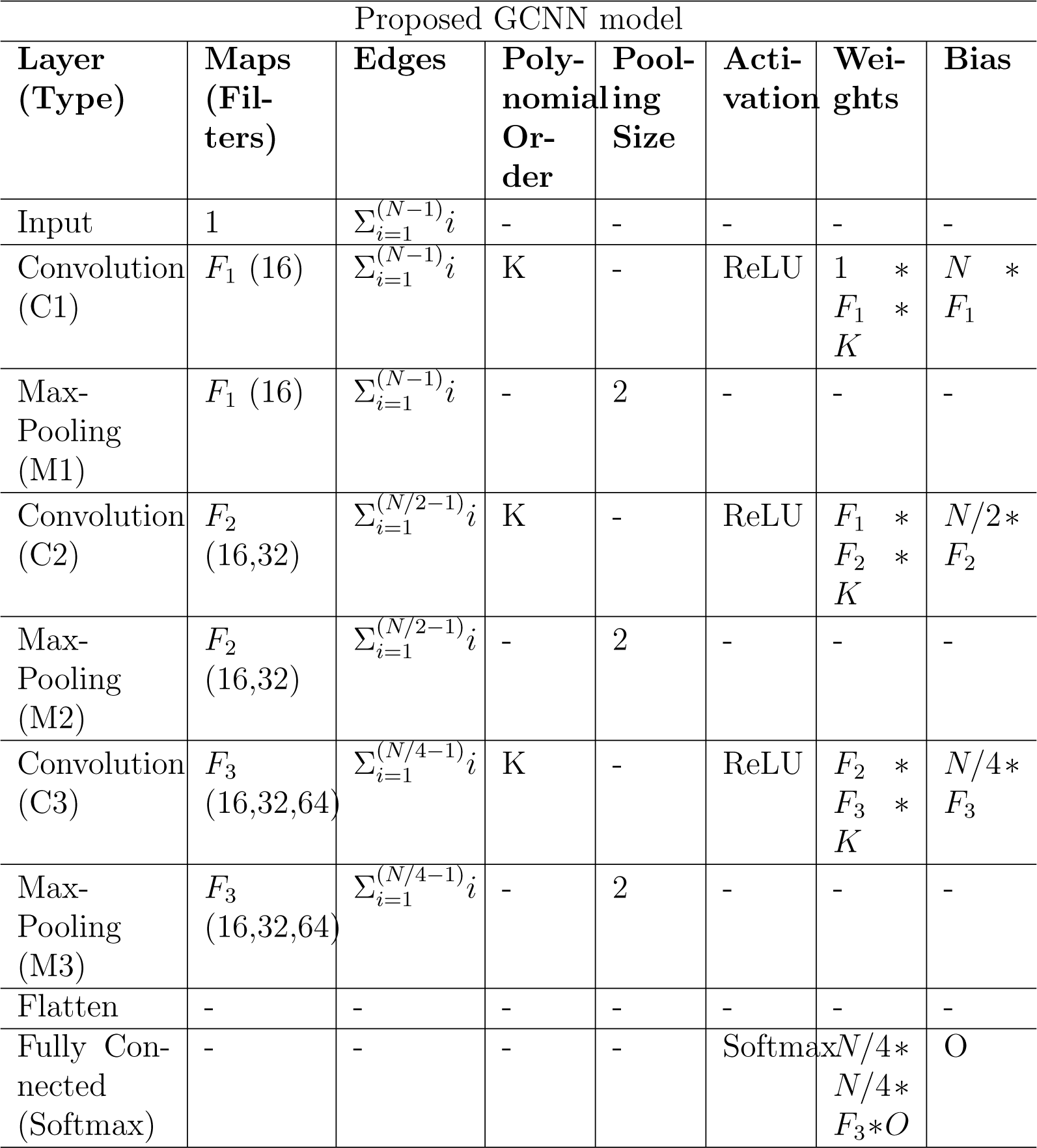
Table shows an implementation of the proposed GCNN model architecture, where O = No. of task, N is = input size, *F_i_ ∈* [*F*_1_*, F*_2_*, F*_3_] = no. of filters at *i_th_* graph convolutional layer, K = polynomial order of filters.

We have used all notations in equations as in previous work [Kipf and Welling, 2016]. For developing Graph Neural Network f(X, A), we considered spatiotemporal connectivity-based multilayer Graph convolutional neural network with equation (3) that indicated forward propagation rule:

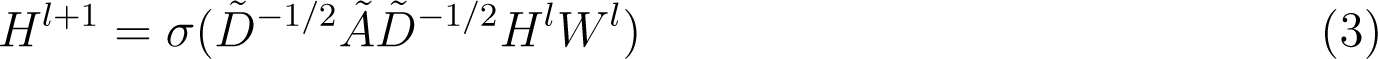

Where *Ã* denotes adjacency matrix i.e. *A* + *I_N_*, whereas layer-wise particular weighted matrix is denoted by *D̃_ii_* = Σ*_j_Ã_ij_* and *W^l^*. ReLU(.) is denoted by *σ*(.) and particularly layer-wise activation matrix is defined by *H^l^* [Kipf and Welling, 2016].

#### 2.3.2 Spectral Graph Convolutional

Following techniques in [Kipf and Welling, 2016], we contemplated spectral convolutions on graphs (GCNs), which are defined as a signal’s multiplication *x ∈ R^N^* by a filter *g_θ_* = *diag*(*θ*) using equation (4). The graph Laplacian’s eigendecomposition in the Fourier domain was calculated via spectral GCNs using the Laplacian matrix.

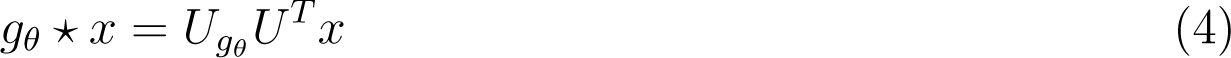

Where *U* = *U*_0_*, U*_1_*, …U_n−_*_1_ denotes eigenvector matrix, and *U^T^ x* denotes transformation from graph Fourier to a signal x [Kipf and Welling, 2016].

Due to the multiplication eigenvector matrix U, which is a complete matrix with n Fourier functions, this procedure is costly. To avoid quadratic complexity, the authors in [Yan et al., 2019] suggested the ChebNet model, which ignores the eigendecomposition by utilizing Laplacian’s learning function. The filter *g_θ_* is estimated via the ChebNet model using Chebyshev polynomials of the diagonal matrix of eigenvalues, as illustrated in equation (5):

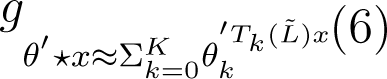

Where diagonal matrix Λ *∈* [*−1*, 1] and Λ̃ = 2Λ*/λ_max_ − I_N_* and Chebyshev polynomial is denoted as *T_k_*(*x*) = 2*xT_k−_*_1_(*x*) *− T_k−_*_2_(*x*) with *T*_0_(*x*) = 1 and *T*_1_(*x*) = *x*.

We calculated convolution of signal x with *g_θ_1* filter using equation (6) [Kipf and Welling, 2016]:

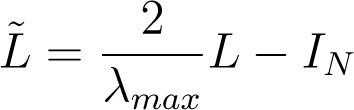

Where *L̃* = ^2^ defines greatest eigenvalue of L.

#### 2.3.3 Training and Testing

We trained our model on SFC matrices and ISFC matrices separately. We divided the data into 80:20 ratios. We used learning rate=0.001, dropout=0.65, and weight decay=0.0, patience =3 [Saeidi et al., 2022]. As batch size is one of the most crucial hyperparameters to modify, a set of batch size values was also considered. This study was implemented using an Adam optimizer with batch sizes of B=[16, 32, 48, 64] across 100 epochs, lowering the learning rate [Saeidi et al., 2022]. For the final prediction, we used the Softmax activation function using equation (7):

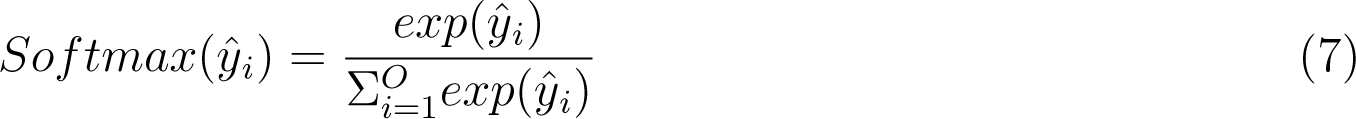

Where, *y*^*_i_ ∈* [*y*^_1_*….y*^*_O_*] = predicted probability of *i_th_* task. Additionally, the optimization function was run using cross-entropy loss using equation (8):

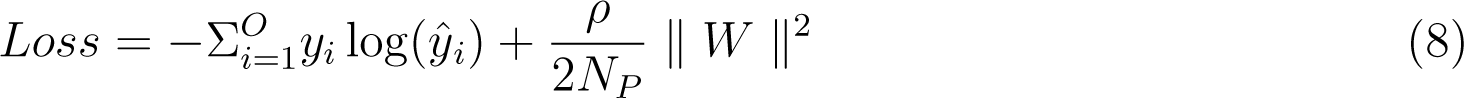

Where *y_i_* = targetted tasks, W = network parameters, *N_P_* = no. of parameters, and *ρ* = weight decay rate. To validate the results, we also implemented five-fold cross-validation.

#### 2.3.4 Identification of Neurobiological Features and Analysis of Result using Five-Fold Cross Validation

The SHAP (SHapley Additive exPlanations) feature diagnostic technique was used to determine the neurological features that contributed most to decoding brain tasks. The SHAP method decomposes the final output prediction to determine the contribution of each attribute to the explainability of machine learning and deep learning models by utilizing equation (9):

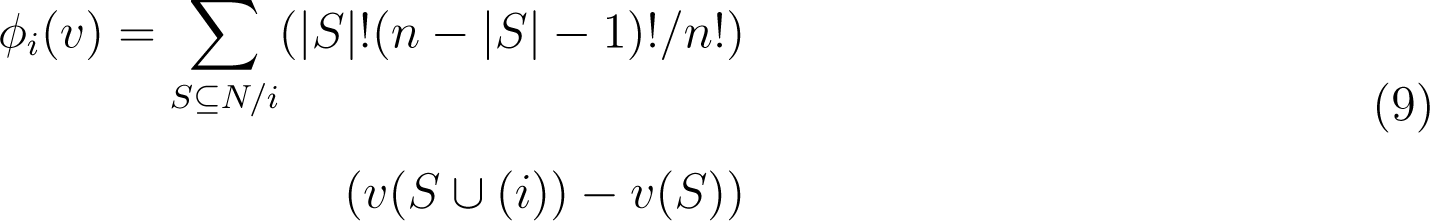

Where: n=total no. of regions, N = all possible subsets, *|S|* no. of features in S, and v(s) calculated contribution of S. To reduce the chance of bias and report low variance, we implemented five-fold cross-validation method to evaluate the model’s performance (precision, recall, accuracy, F1-score).

### 2.4 Prediction of Individual Performance in False-belief Task using Explainable Convolutional Variational Autoencoder Model

In a previous study [Richardson et al., 2018], the authors conducted a ToM-based false-belief task for the 3-12 yrs age group after fMRI scanning and divided all participants into three groups, i.e., pass, fail, and inconsistent, based on their performance. The previous studies [Li et al., 2019a, Li et al., 2019b,Finn and Bandettini, 2021] reported an association between brain signals and behavioral scores in resting state and during movie-watching stimuli. We hypothesized that functional connectivity between brain regions could predict individual performance in false-belief tasks. To check our hypothesis, we considered 122 samples’ age ranges from 3-12 years, which included 84 passes, 15 failures, and 23 inconsistent samples. We conducted this analysis in three ways: a) including all 12 brain regions; b) including dominant brain regions (Total of 8). After decoding brain tasks, we extracted brain regions that contributed the most to prediction. We extracted three dominant regions from ToM networks and three dominant regions from pain networks (six regions from ISFC-based analysis and six regions from SFC-based analysis (there was overlap between the regions, so we considered a total of 8 regions)); c) including only six ToM network regions. We extracted time-series signals and calculated each participant’s static functional connectivity matrix. Finally, we proposed an Explainable Convolutional Variational Auto-Encoder model (Ex-Convolutional VAE), in which we provided SFC matrices of each participant as input and performed prediction of individual performance in false-belief tasks and categorized them into pass, fail, or inconsistent groups.

The proposed Ex-Convolutional VAE model included 2D convolutional layers with ReLU activation function followed by flattening and dense layers with ReLu activation (kernal:3, filters: 32, strides: 2, epoch: 50, latent dimension: 32, no. of channel: 1, batch size: 128 for training Ex-Convolutional VAE and 32 for prediction, padding: SAME, activation function: ReLU for training and sigmoid for prediction) (Refer to Figure 4 and Table 4). The dense layer was used to produce an output of the mean and variance of the latent distribution. Using the reparameterization technique, the sampling function used mean and log variance to sample from latent distribution. The decoder architecture included a dense layer followed by a resampling layer and 2D transposed convolutional layers with ReLU activation function. We used mean squared error (MSE) and Kullback-Leibler (KL) techniques to calculate the loss. The reason for using KL was its ability to regularize learned latent distribution to be close to standard normal distribution. We used trained Ex-Convolutional VAE Latent space for training prediction model with ADAM optimization technique. We performed the prediction using the sigmoid activation function and binary-cross entropy to calculate the loss function (epochs: 50).

**Figure 4:**
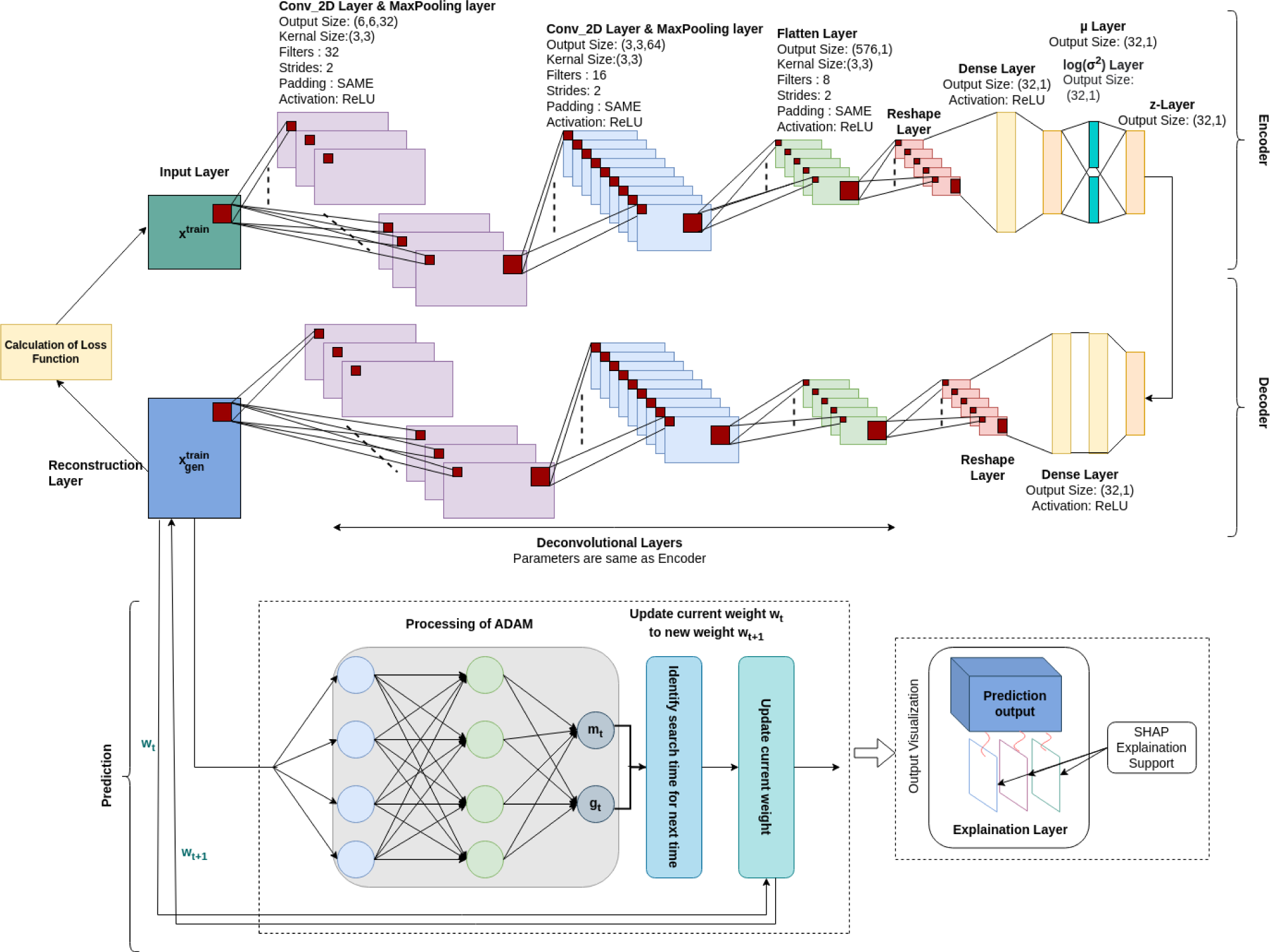
Architecture of proposed Ex-Convolutional VAE model for predicting false-belief task-based pass, fail, and inconsistent groups. The proposed architecture first used the 2D convolutional approach to create a 32-dimensional latent space and trained the prediction model using the ADAM optimization approach (a 32-D latent space was used to train the model). The SHAP approach is used for the explainability of the proposed model.

**Table 4:**
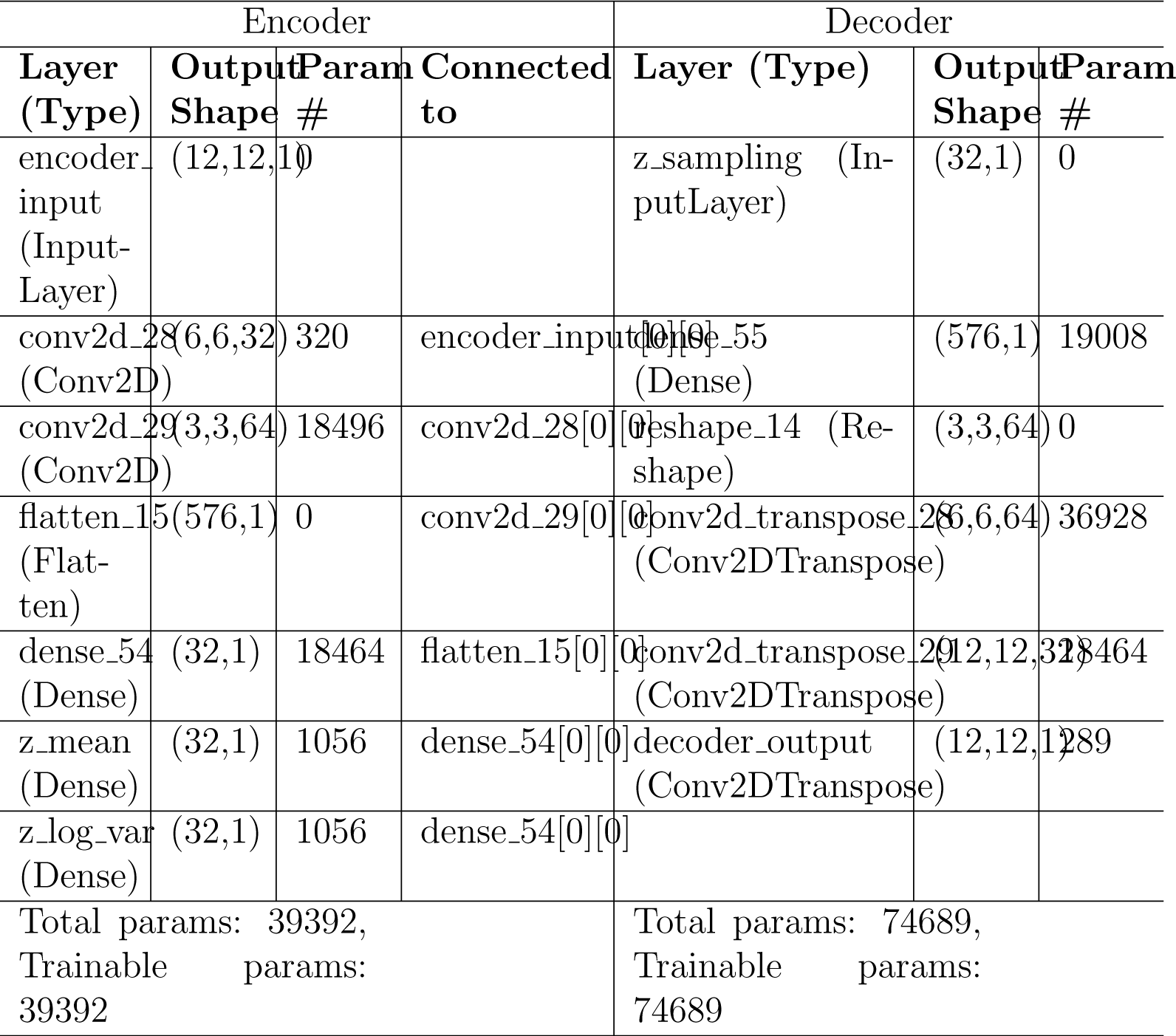
Table shows proposed architecture of Ex-Convolutional VAE model and the parameters with values that have been used on it.

#### 2.4.1 Proposed Approach

In a variational autoencoder model, the encoder produces latent space from a given input while the decoder produces output from this latent space. The decoder inferences that the latent vectors have a normal probability distribution; the parameters of that which are the mean and variance of the vectors, calculated using equation (10) [Lee et al., 2022]:

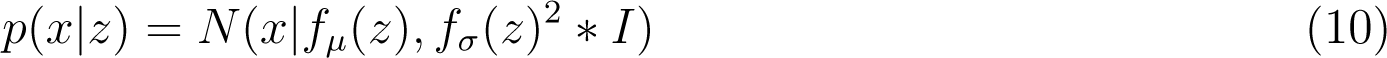

Where *p*(*x|z*) = assumed probability distribution, *f_µ_*(*z*) = log of latent space, and *f_σ_*(*z*)^2^ = variance of latent space. In this particular circumstance, the marginal likelihood estimation technique can be used to the best of its ability to maximize the log-marginal likelihood of the model using equation (11):

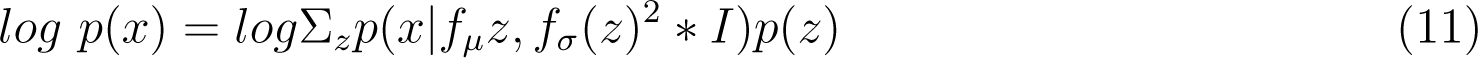

However, it is challenging to maximize the log-marginal likelihood in this form. As a result, we develop variational inference, which simplifies the range of possible outcomes by approximating the posterior probability distribution [Zhang et al., 2018]. An approximately normal probability distribution is an appropriate approximation for the posterior probability distribution. Applying the learning method may be challenging if the input has a high dimension [Lee et al., 2022]. To resolve this, the inferred probability distribution is calculated as a function of x using equations (12) and (13):

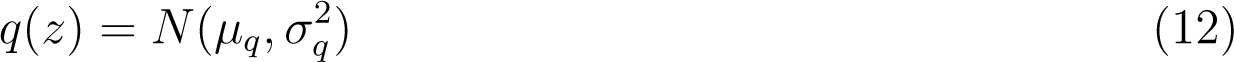

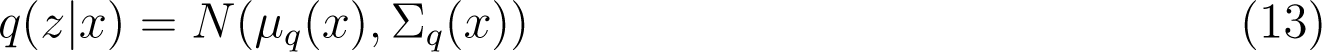

Where *q*(*z*) = inferred normal probability distribution, and *q*(*z|x*) = function of x. Finally, we can obtain the latent vector z by combining the mean value with the product of the inferred normal distribution and the variation. The term ”reparameterization trick” refers to the process used to add a new parameter or feature expressed by equation (14):

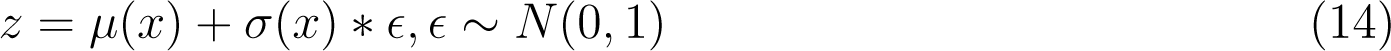

Where z = latent space, and *E* = normally distributed random variable.

The Kullback–Leibler divergence is used to calculate loss function by updating weights and biases that calculate the difference between the actual posterior distribution and inferred distribution [Kingma and Welling, 2013, Kullback and Leibler, 1951] using equation (15):

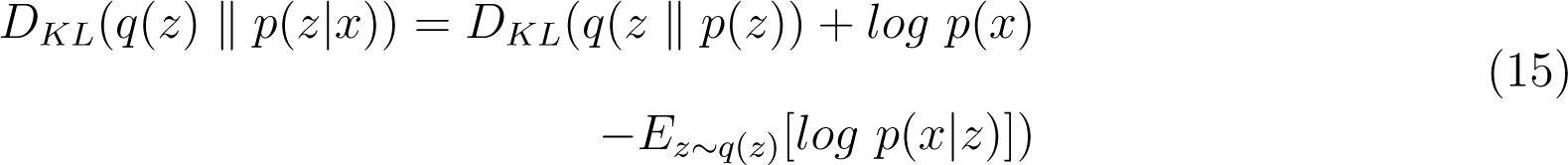

Using the above equation, the log-marginal likelihood of the decoder can be expressed by equation (16):

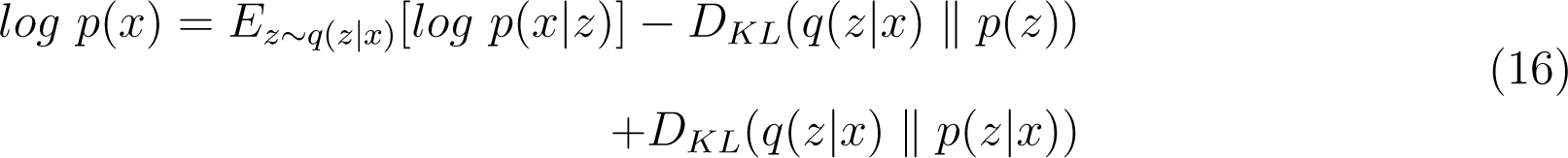

A positive value is always returned by the Kullback–Leibler divergence. As a result, the inequality that results is correct at all times (refer to equation (17)).

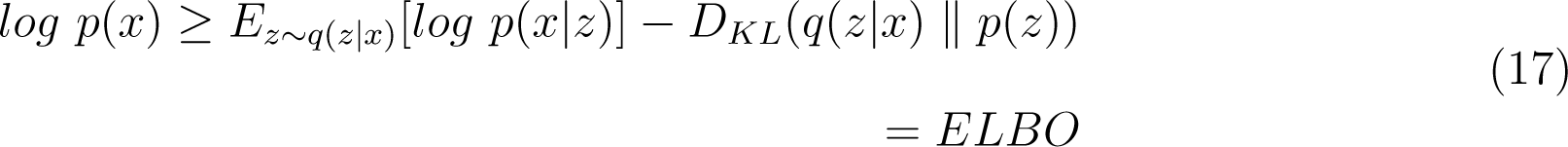

This concept is referred to as effective lower bound (ELBO). Since this inequality is always valid, increasing the ELBO value leads to an increase in the decoder’s log-marginal likelihood. Calculating the loss function of the VAE by multiplying the right-hand side of the equation by a negative value is possible. The following loss function (Refer to equation 18) is used to calculate the training of the convolutional variational autoencoder model:

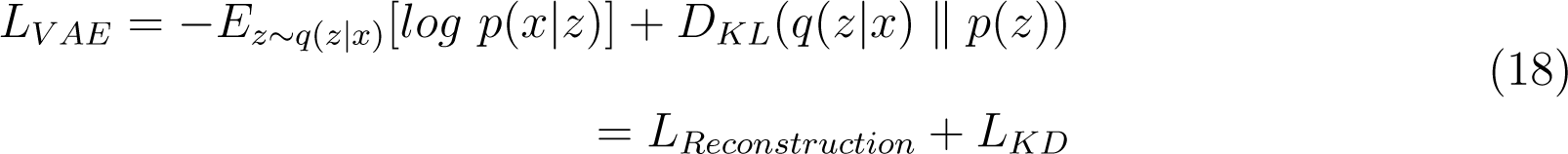

Where *L_Reconstruction_*represents the reconstruction loss, which is the autoencoder’s cross-entropy calculated using input and output data, and *L_KD_* indicates Kullback divergence regularizer value, becomes less as the inferred probability distribution gets closer to a zero-mean Gaussian distribution.

## 3 Results

### 3.1 Computation of SFC and ISFC Matrices

We performed analysis in two ways: a) including the complete dataset, and b) considering age-wise sub-groups, i.e., 3-yrs,4-yrs, 5-yrs, 7-yrs, 8-12 yrs, 3-5 yrs, 7-12 yrs, and adults. The dataset was divided into subgroups to check the effect of age on the model’s performance. Literature informs that [Richardson et al., 2018], networks are not adequately segregated from each other in early childhood. While we considered a dataset that included data from 3-year-old children, it remained a question of whether age is a dependent parameter. We extracted time-series signals from ToM and pain networks for all 12 ROIs (six for ToM, and six for pain) where higher activation occurred. Therefore, SFC matrices of size 12*12 were constructed by calculating Pearson’s correlation for each individual. Additionally, we calculated ISFC matrices of size 12*12 for ToM and pain networks. To validate our results, we performed multiple one-sample t-tests for each connection with a p-value *<* 0.01 and applied FDR correction.

### 3.2 Decoding of Brain Task using Ex-stGCNN model

For decoding tasks or predicting the brain state, we implemented the proposed Ex-stGCNN model. We used SFC and ISFC matrices as separate feature sets to check whether ISFC, a stimulus-driven feature set, could decode tasks better than a static feature set, the SFC. We considered datasets from the early childhood stage, where segregation of each network didn’t happen very well, and activation of other networks like visual networks and default-mode networks was possible. To clarify how the activation of other networks at the same time could affect the model’s performance, we performed an analysis on the whole brain. We compared the results of task decoding using 12 ROIs (ToM and Pain networks) with decoding using the whole brain. We also compared the performance of traditional existing models like MVPA [Haxby et al., 2001], LSTM-RNN [Li and Fan, 2019], and CNN [Wang et al., 2020] and proposed model. We found better results from the proposed model than other existing models (Refer to Table 6).

**Table 6:** Table shows the comparison between performance of traditional models and proposed model on complete dataset.

| Feature Set<br>(Including<br>12 ROIs) | Sr.<br>No. | Models | Accuracy | F1-Score |
| --- | --- | --- | --- | --- |
| **SFC** | 1.a | MVPA | 73.51% | 0.74 |
|  | 1.b | LSTM-RNN | 77.80% | 0.76 |
|  | 1.c | CNN | 79.21% | 0.80 |
|  | 1.d | Ex-stGCNN<br>(Node2Vec) | **85.68%** | **0.84** |
| **ISFC** | 2.a | MVPA | 79.23% | 0.70 |
|  | 2.b | LSTM-RNN | 83.55% | 0.84 |
|  | 2.c | CNN | 85.92% | 0.86 |
|  | 2.d | Ex-stGCNN<br>(Node2Vec) | **94.35%** | **0.95** |

#### 3.2.1 Using SFC Matrices as Feature Set

##### 1. Analysis on Complete Dataset

We divided the data into an 80:20 ratio and trained the model using SFC matrices. We implemented two different node embedding algorithms: Walklets and Node2Vec. Our observations indicated that Node2Vec outperformed Walklets. Using the Node2Vec algorithm for 3D-Convolutional layers, we achieved an average accuracy of 85% with an F1-score of 0.87 for 12 ROIs, while for the whole brain in the same scenario, we achieved 80% accuracy with an F1-score of 0.79. When Walklets were employed for 3d-convolutional layers, we attained an average accuracy of 78% with an F1-score of 0.80 for 12 ROIs and 73% accuracy with an F1-score of 0.72 for the whole brain. However, the model did not perform well with 2D-Convolutional layers, yielding an average accuracy of 75% and an F1-score of 0.76 for 12 ROIs, and 68% accuracy with an F1-score of 0.69 for the whole brain using the Node2Vec algorithm. For Walklets in the same scenario, we obtained an average accuracy of 68% with an F1-score of 0.80 for 12 ROIs, and 65% accuracy with an F1-score of 0.66 for the whole brain. The lowest performance was observed with a 1D-Convolutional layer, resulting in an average accuracy of 60% with an F1-score of 0.61 for 12 ROIs, and an average accuracy of 59% with an F1-score of 0.59 for the whole brain using the Node2Vec algorithm. For Walklets, we achieved 61% accuracy with a 0.62 F1-score for 12 ROIs, and 58% accuracy with a 0.59 F1-score for the whole brain. Our results suggest that GCNN with 3D convolutional layers performs better in decoding brain tasks than 2D or 1D convolutional layers, as indicated in Table 5. We validated our results using five-fold cross-validation and achieved an average accuracy of 75% with an F1-score of 0.76.

**Table 5:** Table shows the performance of proposed Ex-stGCNN model using ISFC and SFC matrices.

| Feature Set | Convolutional Layer | Filters | Walklets (Whole Brain) | Walklets (12 ROIs) | Node2Vec (Whole Brain) | Node2Vec (12 ROIs) |
| --- | --- | --- | --- | --- | --- | --- |
| - | - | - | Accuracy | Accuracy | Accuracy | Accuracy |
| **SFC** | 1D | 16 | 58.15% | 61.50% | 59.71% | 60.25% |
|  | 2D | 16,32 | 65.60% | 68.36% | 68.48% | 75.23% |
|  | 3D | 16,32,64 | 73.85% | 78.72% | 80.92% | 85.68% |
| **ISFC** | 1D | 16 | 63.45% | 65.66% | 65.87% | 69.70% |
|  | 2D | 16,32 | 75.61% | 79.01% | 82.86% | 88.21% |
|  | 3D | 16,32,64 | 82.46% | 92.57% | 85.75% | **94.35%** |

##### 2. Analysis on Age-wise Sub-Groups

Additionally, we analyzed age-wise subgroups to check effect of age on the model’s performance (Refer to Figure 7). We achieved the lowest accuracy of 50% with an F1-score of 0.48 for the 3-yrs age group using Walklets, and the pattern was the same for 4-yrs. We observed a change in the model’s performance from the 7-yrs age group with an accuracy of 68% with an F1-score of 0.69 for 12 ROIs and achieved the highest accuracy for adult groups with an average accuracy of 85% with an F1-score of 0.84. We validated our results using five-fold cross-validation and achieved an average accuracy of 68% with an F1-score of 0.67.

For explainability, we applied SHAP(Shapley Additive exPlanations), which provided the extent to which each input feature contributed to the prediction. We computed the median of feature scores and identified ROIs that contributed the most to classification. We observed that bilateral Temporoparietal Junction (LTPJ and RTPJ), Ventromedial Prefrontal Cortex (vmPFC), Left Interior Insula, and Bilateral Middle Frontal Gyrus (LMFG and RMFG) contributed most to the prediction (Refer to Figure 6).

#### 3.2.2 Using ISFC Matrices as Feature Set

##### 1. Analysis on Complete Dataset

We hypothesized that stimulus-driven measures could better predict the brain state. To test our hypothesis, we calculated ISFC matrices and trained the model. During testing, we achieved the highest accuracy of 94% with an F1-score of 0.95 for 12 ROIs and 85% with an F1-score of 0.87 for the whole brain using Node2Vec for 3D-convolutional layers (Refer to Table 5). In contrast, we obtained an average accuracy of 92% with a 0.93 F1-score for 12 ROIs and 82% accuracy with a 0.83 F1-score for the whole brain using Walklets for 3D-convolutional layers. For 2D-convolutional layers, we achieved 88% with an 0.89 F1-score for 12 ROIs and 82% accuracy with a 0.83 F1-score for the whole brain using Node2Vec. We also obtained 79% accuracy with a 0.79 F1-score for 12 ROIs and 75% accuracy with a 0.76 F1-score for the whole brain using Walklets. Regarding the 1D-convolutional layer, we achieved 69% accuracy with a 0.70 F1-score for 12 ROIs and 65% accuracy with a 0.65 F1-score for the whole brain using Node2Vec. Finally, we achieved 65% accuracy with a 0.66 F1-score for 12 ROIs and 63% accuracy with a 0.64 F1-score for the whole brain using Walklets. Hence, our hypothesis was correct: ISFC measures provided better results compared to FC measures (Refer to Figure 5). To validate the results, we conducted a five-fold cross-validation and achieved an average accuracy of 92% with an F1-score of 0.91.

**Figure 5:**
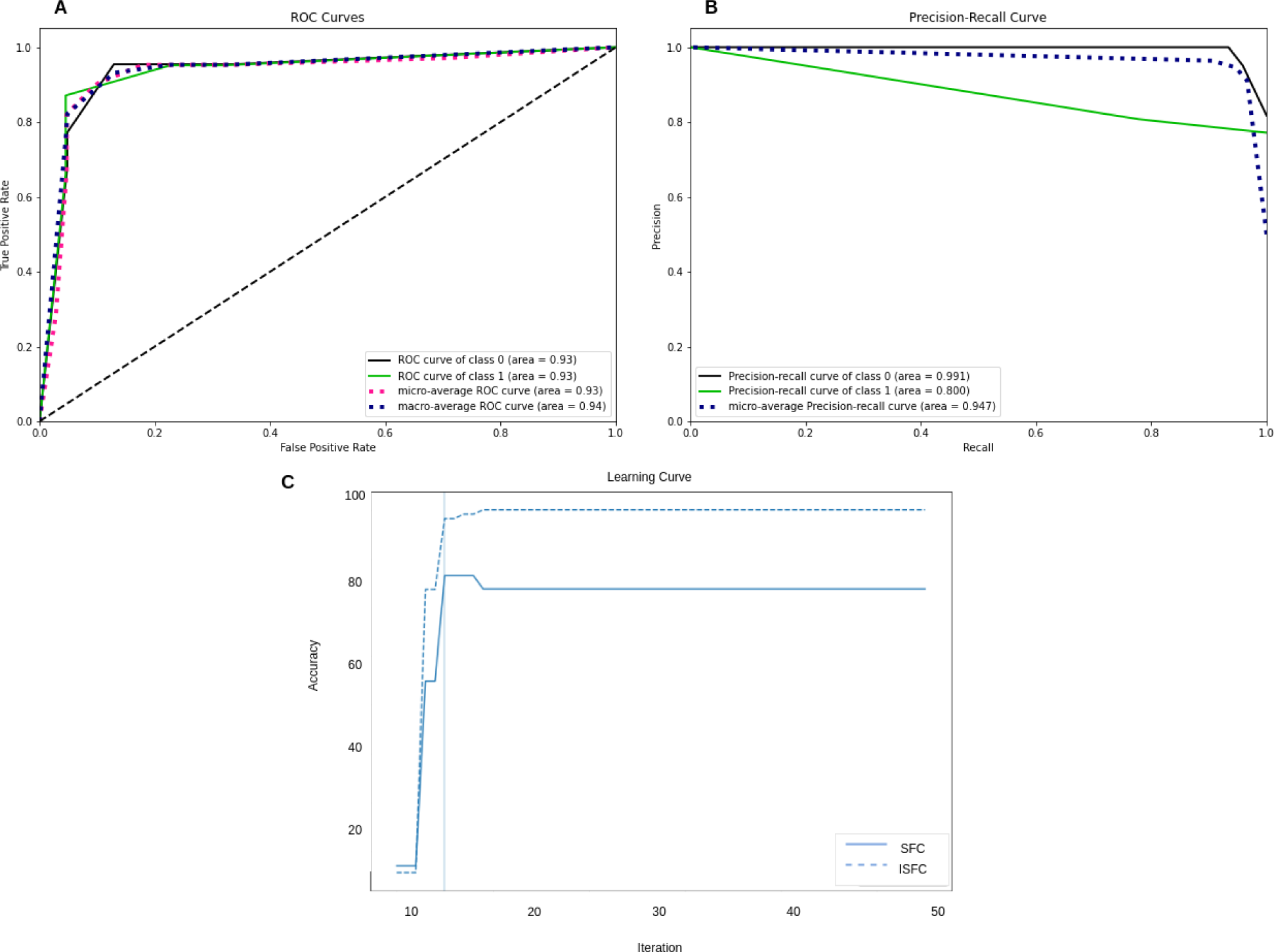
Performance of Ex-stGCNN model: **A and B** show ROC and precision-recall curves of Ex-stGCNN model using ISFC matrices. **C)** shows average accuracy of Ex-stGCNN model using SFC and ISFC matrices separately. Our results indicate that the proposed model gives better results using the ISFC feature set.

##### 2. Analysis on Age-wise Sub-Groups

We analyzed age-wise sub-groups and achieved better results using ISFC matrices. We achieved the best accuracy of 74% with an F1-score of 0.75 for 12 ROIs for the 3-year age group. This proves that even if the segregation of networks didn’t happen appropriately during early development, ISFC measures could predict tasks up to a reasonable threshold (Refer to Figure 7).

Using the SHAP explainability method, we observed that bilateral Temporoparietal Junction (LTPJ and RTPJ), Posterior Cingulate Cortex (PCC), Right Interior Insula, and Bilateral Middle Frontal Gyrus (LMFG and RMFG) contributed most to the prediction of brain state (Refer to Figure 6).

**Figure 6:**
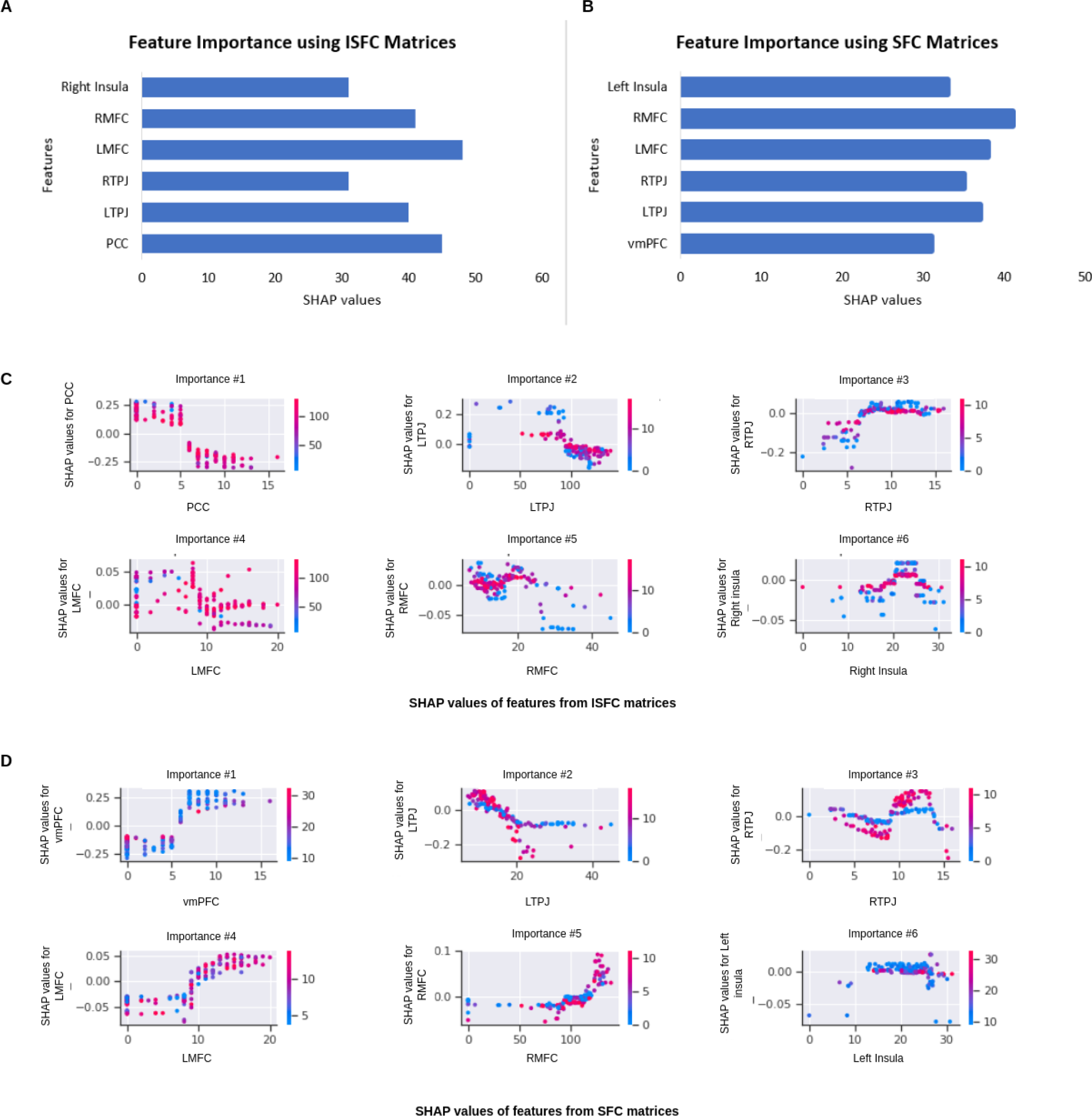
Feature Importance. **A and B** show brain regions that contributed most to the prediction using ISFC and SFC matrices. We extracted three dominating ToM regions and three dominating pain regions. **C and D** show SHAP values of dominating brain regions. There is an overlap between the regions identified by ISFC and SFC matrices.

**Figure 7:**
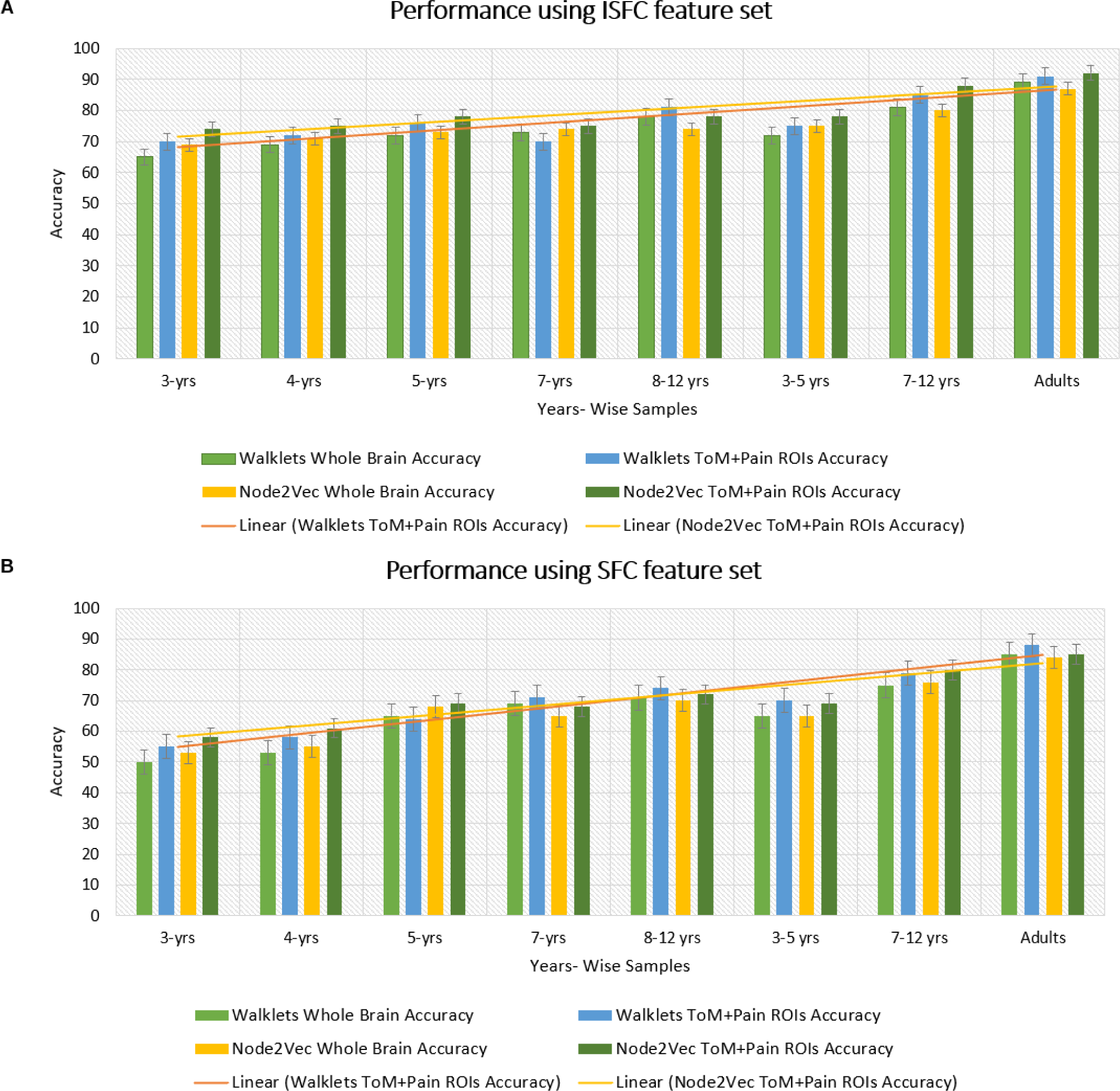
Performance of GCNN model on Age-wise sub-groups: **A)** shows accuracy of proposed Ex-stGCNN model on sub-groups using ISFC matrices. **B)** shows accuracy using SFC matrices. Our results suggest that we can achieve considerable performance using ISFC matrices at early childhood stage. In sub-group analysis, both Walklets and Node2Vec node embedding methods performed approximately the same.

### 3.3 Prediction of Individual Performance on False-belief tasks

In the literature [Li et al., 2019a, Li et al., 2019b, Finn and Bandettini, 2021], an association between brain signals and behavioral scores has been found. We hypothesized that functional connectivity between selected brain regions could predict individual performance on false-belief tasks. To check our hypothesis, we extracted functional connectivity matrices from selected brain regions for each individual. We observed hyper-connectivity for false-belief task-based pass group, whereas moderate connectivity for an inconsistent group and hypo-connectivity for fail group (Refer to Figure 8). To validate our results, we performed multiple one-sample t-tests, one for each connection, with a p-value *<* 0.01, and applied FDR correction.

**Figure 8:**
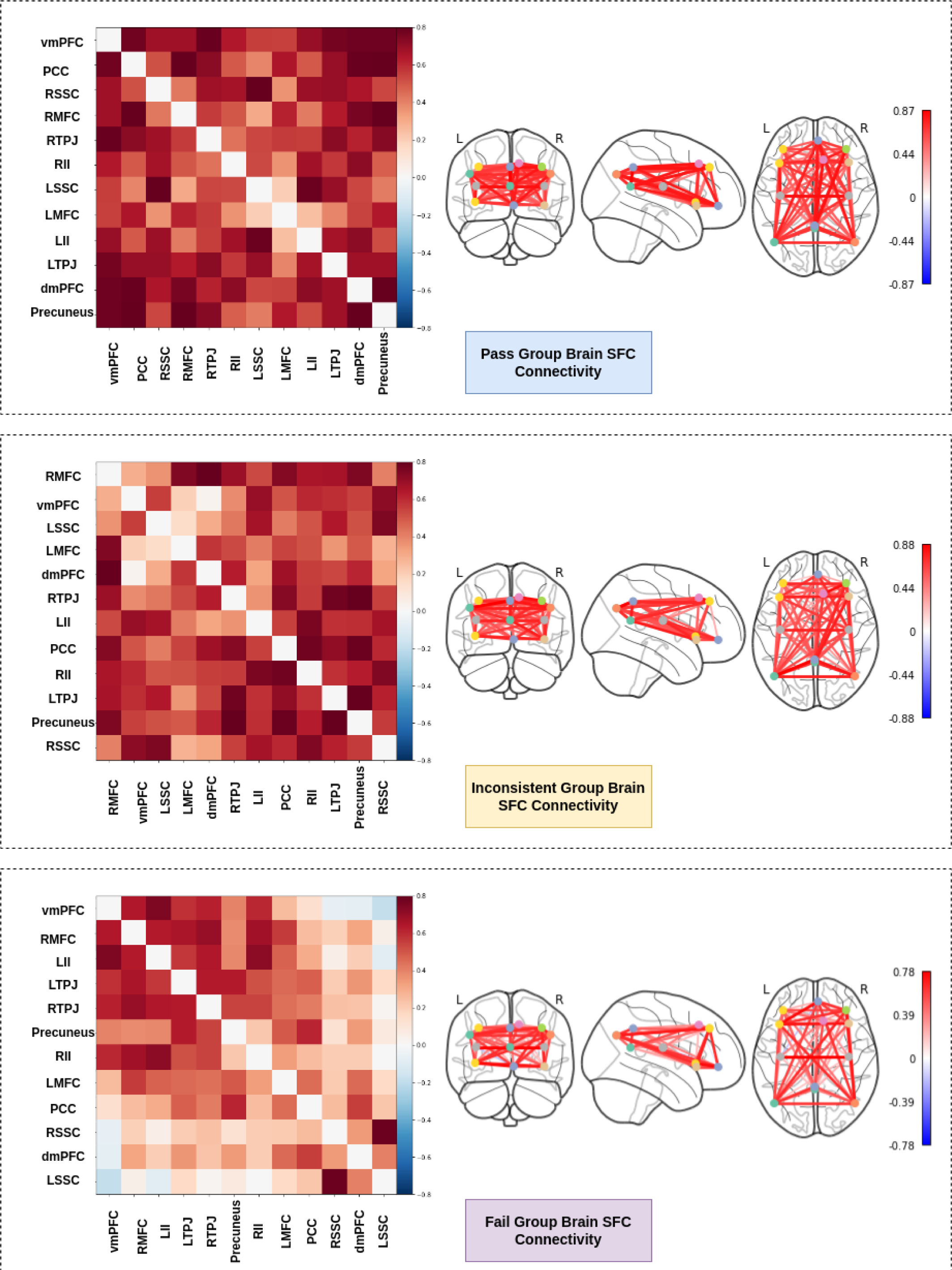
Functional connectivity between ToM _3_a_2_nd pain networks for false-belief task-based pass, fail, and inconsistent groups. Pass group showed hyper-connectivity between ToM and pain network, whereas inconsistent group showed moderate connectivity, and fail group showed hypo-connectivity.

We proposed an Ex-Convolutional VAE model to predict individual performance on false-belief tasks and categorized participants into pass, inconsistent, and fail groups. We trained the proposed model in 3 ways: a) using all 12 ROIs, b) using 8 ROIs that contributed most (dominant ROIs) in decoding brain tasks, and c) using only 6 ToM ROIs. We divided dataset into 80:20 ratios and provided functional connectivity (SFC) matrices as input to train the model. We achieved 90% accuracy with F1-score of 0.91 using 12 ROIs, 84% accuracy with F1-score of 0.83 using eight dominant ROIs, and 80% accuracy with 0.79 F1-score using six ToM ROIs (Refer to Figure 9). To validate our results, we performed five-fold cross-validation and achieved an average accuracy of 87% with F1-score of 0.88. We also tried 1D convolutional and achieved 81% accuracy with 0.80 F1-score using 12 ROIs, 73% with 0.74 F1-score using 8-dominant ROIs, and 66% accuracy with F1-score of 0.67 using six-Tom ROIs. Our results suggest that 2D convolutional is best suited for false-belief task-based group prediction.

**Figure 9:**
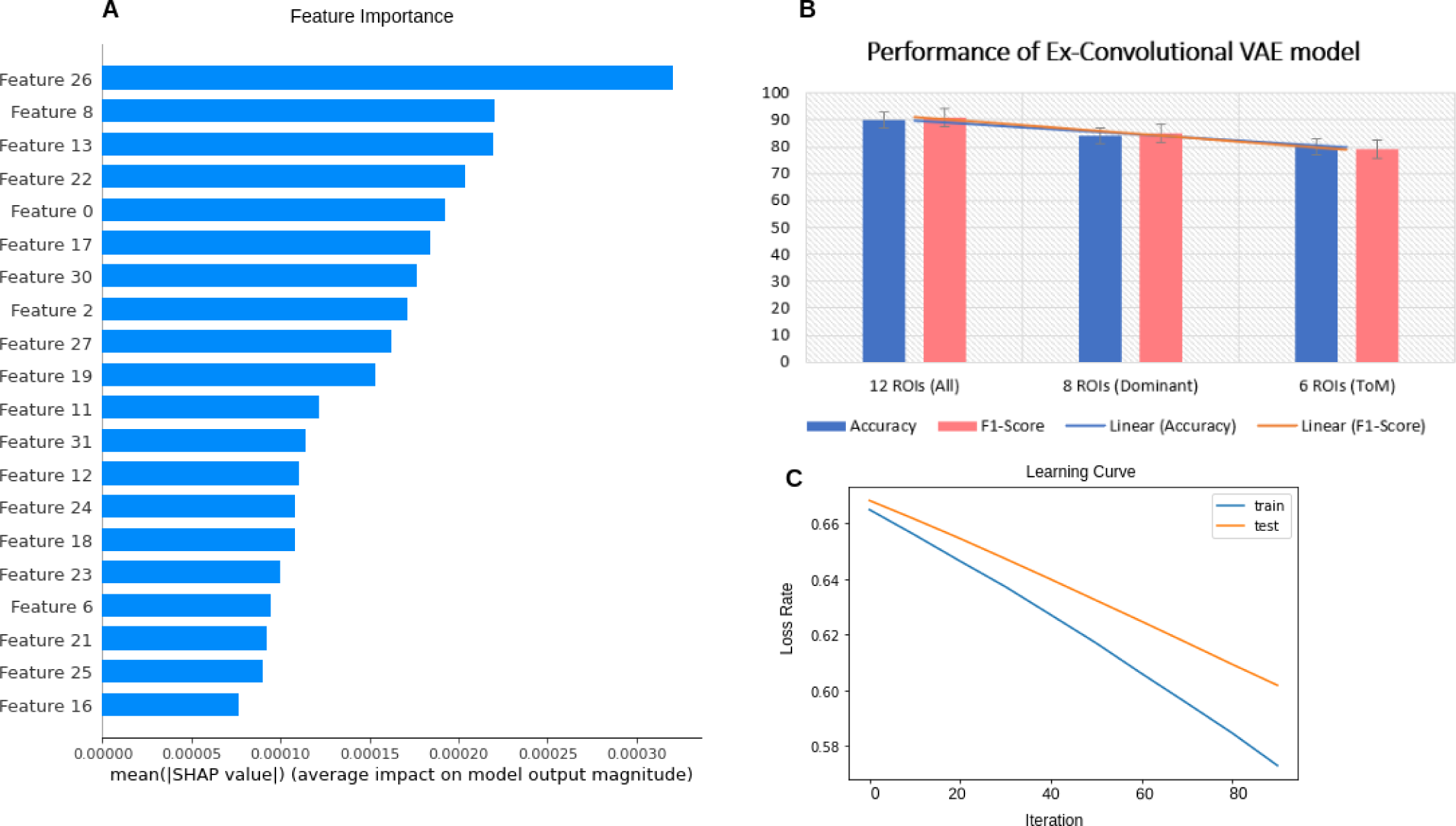
Performance of Ex-Convolutional VAE model: For prediction of groups, the proposed model used 32-dimensional latent space. **A)** Shows importance of features and their SHAP values. **B)** shows performance of proposed model in terms of accuracy and F1-score. **C)** shows the loss rate during training and testing of model.

## 4 Discussion

Decoding brain tasks in the early childhood to pre-adolescent stage using naturalistic stimuli that allow analyzing stimulus-dependent neural processes at different timescales is a challenging issue [Simony and Chang, 2020]. The existing literature [Astington and Edward, 2010, Richardson et al.,

2018] reports that brain networks are not adequately segregated in the early childhood stage (as early as 3-yrs). Young children’s brain development and cognitive abilities undergo substantial transformations during the initial years of their lives [Schult and Wellman, 1997, Schulz et al., 2007, Cohen et al., 2011, Richardson et al., 2018]. Deep learning models showed great success in decoding brain states. Still, existing models [Zhang et al., 2023, Ye et al., 2023, Zhang et al., 2021] suffer from an issue of low accuracy and explainability due to their internal architecture and feature extraction technique. Also, the existing models [Zhang et al., 2023,Ye et al., 2023,Saeidi et al., 2022,Zhang et al., 2021] were tested out in adult data when brain networks are fully matured. Identifying the most effective features that could categorize the relationship between complex naturalistic stimuli and the associated brain activity remains challenging. Moreover, it is pertinent to ask how to design deep learning architecture that could examine the complex representation of developing brain networks.

Firstly, we tested whether a graph convolutional neural network, based on spatio-temporal connectivity, could effectively decode brain tasks in the early childhood developmental dataset using naturalistic stimuli. We believed, since naturalistic stimuli are intrinsically time-bound, a stimulus-driven measure could be a better feature set that captures brain dynamics as compared to a static feature set, which involves a mixture of stimulus-induced neural processes, intrinsic neural processes, and non-neuronal noise to train a deep learning model [Simony and Chang, 2020, Owen et al., 2021]. Secondly, we expected that the functional connectivity between the specified brain regions could predict individual performance in false-belief tasks. In the current study, we proposed an Explainable graph-based deep learning framework to decode brain states (ToM and pain states) and predict individual performance on ToM-related false-belief tasks. We used a dataset in which participants were watching a short ‘Partly Cloudy’ soundless animated movie that was validated to activate ToM and pain networks at particular time points (Refer to Figure 2 and Table 2). The movie used, depicts multiple events that focus on two aspects of the main characters (a cloud named Gus and his stork friend Peck): their bodily sensations (often physical pain) and their mental states (beliefs, desires, and emotions) were salient events that drive developmental change in cortical networks recruited for reasoning about bodies (the pain network) and minds (the theory of mind network). Using a dataset that included developmental age groups (3-12 yrs) and individuals in adulthood opened the opportunity to propose a framework for brain decoding that could analyze complex brain dynamics in the early childhood stage and contextualize these findings from the perspective of adult brains. The movie contained scenes that evoked maximum response magnitude at particular time points in ToM and pain networks. Following the main study of the dataset [Richardson et al., 2018], we selected BOLD signal time-courses from five events (total no. of time-points = 168) belonging to each of the networks at maximum activation (total of ten time-courses)). Using 10 signals from these time-courses, we trained our proposed Ex-stGCNN model to predict whether the participant was experiencing ToM or pain state. We also identified neurobiological brain features as fingerprints that contributed most to the performance. To predict individual performance on false-belief tasks, we proposed an Ex-Convolutional VAE model and categorized participants into pass, fail, and inconsistent groups. We show that our proposed architecture did perform superior to traditional fMRI decoding, RNN, and CNN-based models for complex cognitive states during the naturalistic experience in individuals of early childhood and pre-adolescence, even with short event time courses and small datasets.

### 4.1 Decoding of Brain Tasks using Ex-stGCNN Model

Identifying the relationship between the stimuli and neural activity could be done using decoding of brain states evoked by certain tasks and conditions [Zhang et al., 2023]. Unlike previously proposed methods, i.e., multivariate pattern analysis (MVPA) [Haxby et al., 2001], RNN-based method [Li and Fan, 2019], and CNN-based [Wang et al., 2020] approaches that showed significant results in adult datasets, in this study we proposed a graph-based explainable brain decoding model that combines information on the dynamics of the brain’s distributed networks. Here, we designed an Explainable spatiotemporal Connectivity-based Graph-Convolutional Neural Network (Ex-stGCNN) model to decode brain states that could represent complex topological relationships and interdependencies between data. We selected BOLD time-series data from 10 events (including ToM and pain events) from ToM and pain networks (12 ROIs). We calculated ISFC and SFC matrices and finally trained the model using ISFC and SFC matrices separately to predict whether the participant was experiencing ToM and pain cognitive states (Refer to Figure 3 and Table 3). We achieved 94% accuracy using ISFC matrices involving 12 ROIs, whereas we achieved 85% accuracy using SFC matrices involving 12 ROIs using a 3D convolutional approach. To get more clarity about results, instead of only 12 ROIs, we also analyzed the whole brain SFCs and ISFCs, for the same ten events, but did not get as accurate results. We observed that other networks, like visual network and DMN, were also simultaneously activated, probably which led to the reason we did not get as accurate results in whole brain analysis as with just ToM and pain networks.

We considered a dataset from early childhood, i.e., 3 years, in which network segregation is incomplete. To investigate whether age was a dependent parameter on model performance, we also analyzed age-wise subgroups, i.e., 3-yrs, 4-yrs, 5-yrs, 7-yrs, 8-12 yrs, and adults. We observed that network segregation played an essential role in the model’s performance; we did not achieve considerable performance at age 3-yrs involving the whole brain but achieved better performance using 12 ROIs, and this pattern reminded in 4-yrs individuals (Refer to Figure 7). There was a slight change in the model’s performance from 5 years, and it improved from 5 years to adulthood. We performed analysis using 2D and 1D convolutional approaches but that didn’t give accurate results. We also compared the performance of the proposed model with existing models like MVPA [Haxby et al., 2001], LSTM-RNN [Li and Fan, 2019], and CNN [Wang et al., 2020] and found better results from the proposed model (Refer to Table 6). The unique architecture of the proposed Ex-stGCNN model was able to capture cognitive states even in early childhood stages (as early as 3 years) (Refer to Table 6). The model gave a more accurate prediction for the scenes with high activation. Additionally, we identified neurological brain fingerprints that were responsible for prediction.

Our results suggest that age is a dependent parameter for model performance. The network segregation and activation of other networks could affect the model’s performance at a certain level [Li and Fan, 2018, Wang et al., 2020, Li and Fan, 2019, Gao et al., 2019, Cao et al., 2021, Albouy et al., 2019]. We obtained that even at the early age group, ISFC matrices could capture the activation. We achieved better results using a stimulus-driven feature set i.e., ISFC matrices, which is known to capture the dynamics of social cognition significantly better than a static feature set, i.e., SFC matrices. Following our analysis, we observed that the graph laplacian could be a robust measure for mapping the intrinsic architecture of the human brain that might be accomplished in several ways, including parcellating brain regions, identifying functional areas and networks, and producing connectivity gradients (Refer to Figure 5). This method was effective on static connections in the brain’s connectome and dynamic and relatively transient brain signals that alter over time. Although we got better results using a stimulus-driven feature set, but still got considerable results using a non-stimulus-specific feature set.

### 4.2 Identification of Neurological Brain Finger-prints

Explainability is required to reveal the fundamental decoding process behind the deep learning models that help remove the black-box problem issue. Gradient-based approaches, decomposition methods, and surrogate methods are some techniques developed to explain existing GNNs from various perspectives [Yuan et al., 2022]. In the existing studies, the perturbation-based approaches were implied to identify the link between input characteristics and various outputs [Schlichtkrull et al., 2020, Ying et al., 2019,Yuan et al., 2021]. However, in brain decoding applications, none of these techniques can guarantee the discovery of plausible and comprehensible input characteristics from a neuroscience standpoint.

For the explainability of the proposed model, we implemented the SHAP approach. We identified three dominant brain regions from ToM and three dominant brain regions from pain network for both SFC and ISFC matrices. We determined that bilateral Temporoparietal Junction (LTPJ and RTPJ), Ventromedial Prefrontal Cortex (vmPFC), Left Interior Insula, and Bilateral Middle Frontal Gyrus (LMFG and RMFG) contributed most to the prediction using the SFC feature set (Refer to Figure 6). In contrast, bilateral Temporoparietal Junction (LTPJ and RTPJ), Posterior Cingulate Cortex (PCC), Right Interior Insula, and Bilateral Middle Frontal Gyrus (LMFG and RMFG) contributed most to the prediction using the ISFC feature set.

### 4.3 Prediction of Individual Performance in False-Belief Taks

The previous study [Ganesan et al., 2022] reported that behavioural measures related to visual stimuli could influence the model’s performance in classifying between rest and task states. Given that, we expected that SFC between specified brain regions could effectively predict individual performance in false-belief tasks. To check our hypothesis, we designed an Explainable Convolutional Variational Autoencoder model (Refer to Figure 4 and Table 4). In our analysis, we observed hyper-connectivity for the false belief task-based pass group, moderate connectivity for an inconsistent group, and hypo-connectivity for the fail group (Refer to Figure 8) amongst the 12 selected ROIs. Here, we trained the model using SFC matrices of each individual. We noted that when we used ToM and pain networks, we got better results, i.e., 90% accuracy, whereas using dominant ROIs got 84% accuracy and only using ToM ROIs achieved 80% accuracy (Refer to Figure 9). Our results indicate that pain network, though not as much as ToM, was essential in predicting false-belief task-based groups. However, most failed samples belonged to 3-yrs- and 4-yrs age groups. Given that during this age, the sensory areas are highly activated, and the network segregation doesn’t happen considerably enough as indicated by observed hyper-connectivity between ToM and the pain network, it might be that the proposed model’s learning corresponds to the suboptimal performance on false-belief task by the children of this age. However, we also speculate that this low accuracy might be due to a small sample from this age group.

The proposed work is novel because, to the best of our knowledge, no such framework exists that using a brain-inspired architecture can considerably predict complex brain states and tasks using such short time-course signals from a single fMRI session of early childhood and pre-adolescence individuals. Moreover, the performance of the proposed model to predict the false-belief task indicates the individual’s sensory and social cognition development and opens an avenue for predicting atypical development.

### 4.4 Conclusion and future aspects

The study aimed to propose a framework that can decode higher-order cognitive tasks and associated brain states in short time courses for developmental age group dataset collected from single session recordings without using feature engineering and which can also predict individual performance on false-belief tasks and categorize them in pass, fail, and inconsistent subject groups. We trained the model using ISFC and SFC matrices separately and achieved 94% accuracy using ISFC matrices and 85% using SFC matrices. We also analyzed age-wise subgroups to check the effect of age on the model’s performance. Due to a lack of nuanced network segregation, the model gives lower accuracy for early age groups, i.e., 3 years and 4 years, as for the 5 years and above. We used the SHAP approach to determine the brain fingerprints that contributed most to the prediction. To predict false-belief task-based pass, fail, and inconsistent groups, we proposed an Ex-Convolutional VAE model and achieved 90% accuracy. We validated our results using five-fold cross-validation. The limitations of this work are: a) Dataset size: The dataset only contains 155 samples. Though we showed that our model could still perform well given a small dataset, we believe that a larger sample size, especially compared to the too-small sub-dataset of ages ranging from 3 to 5, might help improve our model performance for this age group (b) primarily healthy participants were considered in this study: to delineate the effects with greater clarity and for better prediction with potential clinical applications age and gender-matched atypical neurodevelopmental samples may be required for a better understanding of the development of these higher-order cognitive networks linked with associated behavior.

### Data Availability

All the analysis codes and implementation of models are available from the GitHub repository (https://github.com/dynamicdip/). All the codes and processed fMRI and behavioral data are available from the corresponding author on reasonable request.

## Acknowledgments

We acknowledge the generous support of IIT Jodhpur Core funds and the Computing facility. D.R. acknowledges the generous support of the NBRC Flagship program BT/MEDIII/NBRC/Flagship/Program/2019: Comparative mapping of common mental disorders (CMD) over the lifespan.

## References

1. Alamolhoda, M., Ayatollahi, S. M. T., and Bagheri, Z. (2017). A comparative study of the impacts of unbalanced sample sizes on the four synthesized methods of meta-analytic structural equation modeling. BMC research notes, 10:1–12.

2. Albouy, P., Caclin, A., Norman-Haignere, S. V., Ĺev^eque, Y., Peretz, I., Tillmann, B., and Zatorre, R. J. (2019). Decoding task-related functional brain imaging data to identify developmental disorders: the case of congenital amusia. Frontiers in Neuroscience, 13:1165.

3. Astington, J. W. and Edward, M. J. (2010). The development of theory of mind in early childhood. Encyclopedia on early childhood development, 14:1–7.

4. Baetens, K., Ma, N., Steen, J., and Van Overwalle, F. (2014). Involvement of the mentalizing network in social and non-social high construal. Social cognitive and affective neuroscience, 9(6):817–824.

5. Bartley, J. E., Boeving, E. R., Riedel, M. C., Bottenhorn, K. L., Salo, T., Eickhoff, S. B., Brewe, E., Sutherland, M. T., and Laird, A. R. (2018). Meta-analytic evidence for a core problem solving network across multiple representational domains. Neuroscience & biobehavioral reviews, 92:318–337.

6. Baucom, L. B., Wedell, D. H., Wang, J., Blitzer, D. N., and Shinkareva, S. V. (2012). Decoding the neural representation of affective states. Neuroimage, 59(1):718–727.

7. Benjamini, Y. and Yekutieli, D. (2001). The control of the false discovery rate in multiple testing under dependency. Annals of statistics, pages 1165–1188.

8. Bhavna, K., Banerjee, R., and Roy, D. (2023a). End-to-end explainable ai: Derived theory-of-mind fingerprints to distinguish between autistic and typically developing and social symptom severity. bioRxiv, pages 2023–01.

9. Bhavna, K., Ghosh, N., Banerjee, R., and Roy, D. (2023b). Developmental stability and segregation of theory of mind and pain networks carry distinct temporal signatures during naturalistic viewing. bioRxiv, pages 2023–08.

10. Burgund, E. D., Kang, H. C., Kelly, J. E., Buckner, R. L., Snyder, A. Z., Petersen, S. E., and Schlaggar, B. L. (2002). The feasibility of a common stereotactic space for children and adults in fmri studies of development. Neuroimage, 17(1):184–200.

11. Bzdok, D., Varoquaux, G., Grisel, O., Eickenberg, M., Poupon, C., and Thirion, B. (2016). Formal models of the network co-occurrence underlying mental operations. PLoS computational biology, 12(6):e1004994.

12. Cantlon, J. F., Brannon, E. M., Carter, E. J., and Pelphrey, K. A. (2006). Functional imaging of numerical processing in adults and 4-y-old children. PLoS biology, 4(5):e125.

13. Cao, L., Huang, D., Zhang, Y., Jiang, X., and Chen, Y. (2021). Brain decoding using fnirs. In Proceedings of the AAAI Conference on Artificial Intelligence, volume 35, pages 12602–12611.

14. Christ, M., Kempa-Liehr, A. W., and Feindt, M. (2016). Distributed and parallel time series feature extraction for industrial big data applications. arXiv preprint arXiv:1610.07717.

15. Cohen, E., Burdett, E., Knight, N., and Barrett, J. (2011). Cross-cultural similarities and differences in person-body reasoning: Experimental evidence from the united kingdom and brazilian amazon. Cognitive science, 35(7):1282–1304.

16. Correia, J., Formisano, E., Valente, G., Hausfeld, L., Jansma, B., and Bonte, M. (2014). Brain-based translation: fmri decoding of spoken words in bilinguals reveals language-independent semantic representations in anterior temporal lobe. Journal of Neuroscience, 34(1):332–338.

17. Demirtaş, M., Ponce-Alvarez, A., Gilson, M., Hagmann, P., Mantini, D., Betti, V., Romani, G. L., Friston, K., Corbetta, M., and Deco, G. (2019). Distinct modes of functional connectivity induced by movie-watching. NeuroImage, 184:335–348.

18. Dubben, H.-H. and Beck-Bornholdt, H.-P. (2005). Systematic review of publication bias in studies on publication bias. Bmj, 331(7514):433–434.

19. Fey, M. and Lenssen, J. E. (2019). Fast graph representation learning with pytorch geometric. arXiv preprint arXiv:1903.02428.

20. Finn, E. S. and Bandettini, P. A. (2021). Movie-watching outperforms rest for functional connectivity-based prediction of behavior. NeuroImage, 235:117963.

21. Ganesan, S., Lv, J., and Zalesky, A. (2022). Multi-timepoint pattern analysis: Influence of personality and behavior on decoding context-dependent brain connectivity dynamics. Human brain mapping, 43(4):1403–1418.

22. Gao, Y., Zhang, Y., Wang, H., Guo, X., and Zhang, J. (2019). Decoding behavior tasks from brain activity using deep transfer learning. IEEE Access, 7:43222–43232.

23. Grover, A. and Leskovec, J. (2016). node2vec: Scalable feature learning for networks. In Proceedings of the 22nd ACM SIGKDD international conference on Knowledge discovery and data mining, pages 855–864.

24. Hasson, U., Nir, Y., Levy, I., Fuhrmann, G., and Malach, R. (2004). Intersubject synchronization of cortical activity during natural vision. science, 303(5664):1634–1640.

25. Haxby, J. V., Connolly, A. C., and Guntupalli, J. S. (2014). Decoding neural representational spaces using multivariate pattern analysis. Annual review of neuroscience, 37:435– 456.

26. Haxby, J. V., Gobbini, M. I., Furey, M. L., Ishai, A., Schouten, J. L., and Pietrini, P. (2001). Distributed and overlapping representations of faces and objects in ventral temporal cortex. Science, 293(5539):2425–2430.

27. Hou, Y., Jia, S., Lun, X., Hao, Z., Shi, Y., Li, Y., Zeng, R., and Lv, J. (2022). Gcns-net: a graph convolutional neural network approach for decoding time-resolved eeg motor imagery signals. IEEE Transactions on Neural Networks and Learning Systems.

28. Huth, A. G., Nishimoto, S., Vu, A. T., and Gallant, J. L. (2012). A continuous semantic space describes the representation of thousands of object and action categories across the human brain. Neuron, 76(6):1210–1224.

29. Jacoby, N., Bruneau, E., Koster-Hale, J., and Saxe, R. (2016). Localizing pain matrix and theory of mind networks with both verbal and non-verbal stimuli. Neuroimage, 126:39– 48.

30. Keil, B., Alagappan, V., Mareyam, A., McNab, J. A., Fujimoto, K., Tountcheva, V., Triantafyllou, C., Dilks, D. D., Kanwisher, N., Lin, W., et al. (2011). Size-optimized 32-channel brain arrays for 3 t pediatric imaging. Magnetic Resonance in Medicine, 66(6):1777–1787.

31. Kim, D., Kay, K., Shulman, G. L., and Corbetta, M. (2018). A new modular brain organization of the bold signal during natural vision. Cerebral Cortex, 28(9):3065–3081.

32. Kingma, D. P. and Welling, M. (2013). Auto-encoding variational bayes. arXiv preprint arXiv:1312.6114.

33. Kipf, T. N. and Welling, M. (2016). Semi-supervised classification with graph convolutional networks. arXiv preprint arXiv:1609.02907.

34. Kullback, S. and Leibler, R. A. (1951). On information and sufficiency. The annals of mathematical statistics, 22(1):79–86.

35. Kumar, M., Anderson, M. J., Antony, J. W., Baldassano, C., Brooks, P. P., Cai, M. B., Chen, P.-H. C., Ellis, C. T., Henselman-Petrusek, G., Huberdeau, D., et al. (2020). Brainiak: The brain imaging analysis kit.

36. Lee, S. M., Park, S.-Y., and Choi, B.-H. (2022). Application of domain-adaptive convolutional variational autoencoder for stress-state prediction. Knowledge-Based Systems, 248:108827.

37. Li, H. and Fan, Y. (2018). Brain decoding from functional mri using long short-term memory recurrent neural networks. In Medical Image Computing and Computer Assisted Intervention–MICCAI 2018: 21st International Conference, Granada, Spain, September 16-20, 2018, Proceedings, Part III 11, pages 320–328. Springer.

38. Li, H. and Fan, Y. (2019). Interpretable, highly accurate brain decoding of subtly distinct brain states from functional mri using intrinsic functional networks and long short-term memory recurrent neural networks. NeuroImage, 202:116059.

39. Li, J., Bolt, T., Bzdok, D., Nomi, J. S., Yeo, B. T., Spreng, R. N., and Uddin, L. Q. (2019a). Topography and behavioral relevance of the global signal in the human brain. Scientific reports, 9(1):14286.

40. Li, J., Kong, R., Líegeois, R., Orban, C., Tan, Y., Sun, N., Holmes, A. J., Sabuncu, M. R., Ge, T., and Yeo, B. T. (2019b). Global signal regression strengthens association between resting-state functional connectivity and behavior. NeuroImage, 196:126–141.

41. Li, W., Wang, Z., Zhang, L., Qiao, L., and Shen, D. (2017). Remodeling pearson’s correlation for functional brain network estimation and autism spectrum disorder identification. Frontiers in neuroinformatics, 11:55.

42. Lieberman, M. D., Burns, S. M., Torre, J. B., and Eisenberger, N. I. (2016). Reply to wager et al.: Pain and the dacc: The importance of hit rate-adjusted effects and posterior probabilities with fair priors. Proceedings of the National Academy of Sciences, 113(18):E2476– E2479.

43. Lieberman, M. D. and Eisenberger, N. I. (2015). The dorsal anterior cingulate cortex is selective for pain: Results from large-scale reverse inference. Proceedings of the National Academy of Sciences, 112(49):15250–15255.

44. Lin, L. (2018). Bias caused by sampling error in meta-analysis with small sample sizes. PloS one, 13(9):e0204056.

45. Lynch, L. K., Lu, K.-H., Wen, H., Zhang, Y., Saykin, A. J., and Liu, Z. (2018). Task-evoked functional connectivity does not explain functional connectivity differences between rest and task conditions. Human brain mapping, 39(12):4939–4948.

46. Mazziotta, J., Toga, A., Evans, A., Fox, P., Lancaster, J., Zilles, K., Woods, R., Paus, T., Simpson, G., Pike, B., et al. (2001). A probabilistic atlas and reference system for the human brain: International consortium for brain mapping (icbm). Philosophical Transactions of the Royal Society of London. Series B: Biological Sciences, 356(1412):1293–1322.

47. Mazziotta, J. C., Toga, A. W., Evans, A., Fox, P., Lancaster, J., et al. (1995). A probabilistic atlas of the human brain: theory and rationale for its development. Neuroimage, 2(2):89–101.

48. Nastase, S. A., Gazzola, V., Hasson, U., and Keysers, C. (2019). Measuring shared responses across subjects using intersubject correlation.

49. Owen, L. L., Chang, T. H., and Manning, J. R. (2021). High-level cognition during story listening is reflected in high-order dynamic correlations in neural activity patterns. Nature Communications, 12(1):5728.

50. Paszke, A., Gross, S., Massa, F., Lerer, A., Bradbury, J., Chanan, G., Killeen, T., Lin, Z., Gimelshein, N., Antiga, L., et al. (2019). Pytorch: An imperative style, high-performance deep learning library. Advances in neural information processing systems, 32.

51. Penny, W. D., Friston, K. J., Ashburner, J. T., Kiebel, S. J., and Nichols, T. E. (2011). Statistical parametric mapping: the analysis of functional brain images. Elsevier.

52. Perozzi, B., Kulkarni, V., Chen, H., and Skiena, S. (2017). Don’t walk, skip! online learning of multi-scale network embeddings. In Proceedings of the 2017 IEEE/ACM International Conference on Advances in Social Networks Analysis and Mining 2017, pages 258–265.

53. Pilgramm, S., de Haas, B., Helm, F., Zentgraf, K., Stark, R., Munzert, J., and Krüger, B. (2016). Motor imagery of hand actions: Decoding the content of motor imagery from brain activity in frontal and parietal motor areas. Human brain mapping, 37(1):81–93.

54. Poldrack, R. A. (2006). Can cognitive processes be inferred from neuroimaging data? Trends in cognitive sciences, 10(2):59–63.

55. Poldrack, R. A. (2011). Inferring mental states from neuroimaging data: from reverse inference to large-scale decoding. Neuron, 72(5):692–697.

56. Poldrack, R. A., Halchenko, Y. O., and Hanson, S. J. (2009). Decoding the large-scale structure of brain function by classifying mental states across individuals. Psychological science, 20(11):1364–1372.

57. Reher, K. and Sohn, P. (2009). Partly cloudy. Motion Picture](Pixar Animation Studios and Walt Disney Pictures, 2009).

58. Richardson, H., Lisandrelli, G., Riobueno-Naylor, A., and Saxe, R. (2018). Development of the social brain from age three to twelve years. Nature communications, 9(1):1–12.

59. Rosenbaum, R., Smith, M. A., Kohn, A., Rubin, J. E., and Doiron, B. (2017). The spatial structure of correlated neuronal variability. Nature neuroscience, 20(1):107– 114.

60. Saeidi, M., Karwowski, W., Farahani, F. V., Fiok, K., Hancock, P., Sawyer, B. D., Christov-Moore, L., and Douglas, P. K. (2022). Decoding task-based fmri data with graph neural networks, considering individual differences. Brain Sciences, 12(8):1094.

61. Santhanam, G., Ryu, S. I., Yu, B. M., Afshar, A., and Shenoy, K. V. (2006). A high-performance brain–computer interface. nature, 442(7099):195–198.

62. Schlichtkrull, M. S., De Cao, N., and Titov, I. (2020). Interpreting graph neural networks for nlp with differentiable edge masking. arXiv preprint arXiv:2010.00577.

63. Schult, C. A. and Wellman, H. M. (1997). Explaining human movements and actions: Children’s understanding of the limits of psychological explanation. Cognition, 62(3):291–324.

64. Schulz, L. E., Bonawitz, E. B., and Griffiths, T. L. (2007). Can being scared cause tummy aches? naive theories, ambiguous evidence, and preschoolers’ causal inferences. Developmental psychology, 43(5):1124.

65. Simony, E. and Chang, C. (2020). Analysis of stimulus-induced brain dynamics during naturalistic paradigms. NeuroImage, 216:116461.

66. Simony, E., Honey, C. J., Chen, J., Lositsky, O., Yeshurun, Y., Wiesel, A., and Hasson, U. (2016). Dynamic reconfiguration of the default mode network during narrative comprehension. Nature communications, 7(1):12141.

67. Song, S., Ma, X., Zhan, Y., Zhan, Z., Yao, L., and Zhang, J. (2013). Bayesian reconstruction of multiscale local contrast images from brain activity. Journal of neuroscience methods, 220(1):39–45.

68. Spellman, T., Rigotti, M., Ahmari, S. E., Fusi, S., Gogos, J. A., and Gordon, J. A. (2015). Hippocampal–prefrontal input supports spatial encoding in working memory. Nature, 522(7556):309–314.

69. Varoquaux, G., Schwartz, Y., Poldrack, R. A., Gauthier, B., Bzdok, D., Poline, J.-B., and Thirion, B. (2018). Atlases of cognition with large-scale human brain mapping. PLoS computational biology, 14(11):e1006565.

70. Wager, T. D., Atlas, L. Y., Botvinick, M. M., Chang, L. J., Coghill, R. C., Davis, K. D., Iannetti, G. D., Poldrack, R. A., Shackman, A. J., and Yarkoni, T. (2016). Pain in the acc? Proceedings of the National Academy of Sciences, 113(18):E2474–E2475.

71. Wang, X., Liang, X., Jiang, Z., Nguchu, B. A., Zhou, Y., Wang, Y., Wang, H., Li, Y., Zhu, Y., Wu, F., et al. (2020). Decoding and mapping task states of the human brain via deep learning. Human brain mapping, 41(6):1505–1519.

72. Whitfield-Gabrieli, S., Nieto-Castanon, A., and Ghosh, S. (2011). Artifact detection tools (art). Cambridge, MA. Release Version, 7(19):11.

73. Xie, H. and Redcay, E. (2022). A tale of two connectivities: intra-and intersubject functional connectivity jointly enable better prediction of social abilities. bioRxiv, pages 2022–02.

74. Yan, Z., Youyong, K., Jiasong, W., Coatrieux, G., and Huazhong, S. (2019). Brain tissue segmentation based on graph convolutional networks. In 2019 IEEE International Conference on Image Processing (ICIP), pages 1470–1474. IEEE.

75. Ye, Z., Qu, Y., Liang, Z., Wang, M., and Liu, Q. (2022). Explainable fmri-based brain decoding via spatial temporal-pyramid graph convolutional network. arXiv preprint arXiv:2210.05713.

76. Ye, Z., Qu, Y., Liang, Z., Wang, M., and Liu, Q. (2023). Explainable fmri-based brain decoding via spatial temporal-pyramid graph convolutional network. Human Brain Mapping, 44(7):2921–2935.

77. Ying, Z., Bourgeois, D., You, J., Zitnik, M., and Leskovec, J. (2019). Gnnexplainer: Generating explanations for graph neural networks. Advances in neural information processing systems, 32.

78. Yuan, H., Yu, H., Gui, S., and Ji, S. (2022). Explainability in graph neural networks: A taxonomic survey. IEEE transactions on pattern analysis and machine intelligence, 45(5):5782–5799.

79. Yuan, H., Yu, H., Wang, J., Li, K., and Ji, S. (2021). On explainability of graph neural networks via subgraph explorations. In International conference on machine learning, pages 12241–12252. PMLR.

80. Zhang, C., Bütepage, J., Kjellström, H., and Mandt, S. (2018). Advances in variational inference. IEEE transactions on pattern analysis and machine intelligence, 41(8):2008– 2026.

81. Zhang, Y. and Bellec, P. (2019). Functional annotation of human cognitive states using graph convolution networks. In Real Neurons {\&} Hidden Units: Future directions at the intersection of neuroscience and artificial intelligence@ NeurIPS 2019.

82. Zhang, Y., Gao, Y., Xu, J., Zhao, G., Shi, L., and Kong, L. (2023). Unsupervised joint domain adaptation for decoding brain cognitive states from tfmri images. IEEE Journal of Biomedical and Health Informatics.

83. Zhang, Y., Tetrel, L., Thirion, B., and Bellec, P. (2021). Functional annotation of human cognitive states using deep graph convolution. NeuroImage, 231:117847.

